# Chemical screening pipeline for identification of specific plant autophagy modulators

**DOI:** 10.1101/569327

**Authors:** Adrian N. Dauphinee, Catarina Cardoso, Kerstin Dalman, Jonas A. Ohlsson, Stina Berglund Fick, Stephanie Robert, Glenn R. Hicks, Peter V. Bozhkov, Elena A. Minina

## Abstract

Autophagy is a major catabolic process in eukaryotes with a key role in homeostasis, programmed cell death and aging. In plants, autophagy is also known to regulate agriculturally important traits such as stress resistance, longevity, vegetative biomass and seed yield. Despite its significance, there is still a shortage of reliable tools modulating plant autophagy. Here we describe the first robust pipeline for identification of specific plant autophagy-modulating compounds. Our screening protocol comprises four phases: (i) high-throughput screening of chemical compounds in cell cultures of *Nicotiana tabacum*; (ii) confirmation of the identified hits *in planta* using *Arabidopsis thaliana*; (iii) further characterization of the effect using conventional molecular biology methods; (iv) verification of chemical specificity on autophagy *in planta*. The methods detailed here streamline identification of specific plant autophagy modulators and aid in unravelling the molecular mechanisms of plant autophagy.

## Main

Autophagy is a key regulator of cellular homeostasis and plays an essential role in abiotic and biotic stress responses, fecundity, aging and cell death^1–6^. The autophagy pathway comprises selective or bulk sequestration of cytosolic contents into double-membraned vesicles known as autophagosomes and their delivery to the lytic compartment, where autophagy activity (or autophagic flux) completes with the degradation of the delivered cargo^7^. Autophagy is coordinated by evolutionary conserved autophagy-related (ATG) proteins that orchestrate sequestration of the cargo, formation and trafficking of the autophagosomes and cargo degradation. Among these proteins, ATG8 is remarkable in the sense that it is incorporated into the membranes of forming autophagosomes, trafficked to the lytic compartment and degraded together with the cargo, making it an ideal tool for autophagy activity read-out^8^.

The importance of autophagy for plant stress tolerance in general and crop performance in particular is of critical interest in plant biology, requiring continuous development of increasingly sophisticated tools for its investigation^1,9^. The forward and reverse genetic approaches such as knockouts, knockdowns and overexpression of *ATG* genes, although being effective tools, have limitations^1,8^. The chemical modulation of autophagy presents an alternative option and has also been instrumental for our current understanding of the underlying mechanisms controlling autophagy^8^. However, an increasing number of studies describe significant off-target effects of commonly used autophagy-modulating compounds. For example, 3-methyladenine (3-MA) that is typically used as an inhibitor of autophagy^8^, can also increase autophagic flux after prolonged treatments due to its transient suppression of class III phosphoinositide 3-kinases (PI3Ks)^10^. Another autophagy inhibitor, chloroquine, has been shown recently to induce autophagy-independent severe disorganization of the Golgi and endo-lysosomal systems^11^. An enhancer of autophagy, rapamycin, extensively used for yeast and animal model systems is known to specifically form a ternary complex with the FKBP12 (12-kDa cis-trans peptidyl-prolyl isomerase FK506 Binding Protein) and the TOR (Target of Rapamycin) kinase^8,12^ to induce autophagy. Nevertheless, an opposite effect is caused by high concentrations of rapamycin^13^. Furthermore, although rapamycin was originally thought to be ineffective in *Arabidopsis thaliana*^14^, potentially due to the low capacity of plant FKB12 proteins to form the complex, a more recent report suggests this is not the case^15^. These studies illustrate the importance and need for precise and robust screening strategies to identify novel autophagy modulators. Furthermore, novel modulators of autophagy may play a critical role in understanding the underlying mechanisms regulating the process.

Here we describe the first robust pipeline for identification of chemical compounds specifically modulating plant autophagy. The assays utilize a combination of reporter and control transgenic lines to optimize the treatment conditions for cell cultures and *in planta* to ensure that the compounds influence autophagic flux without causing considerable stress to other endomembrane compartments. Using this new method, we have identified several promising autophagy-modulating compounds.

## Results

### Screen overview

The pipeline comprises a combination of assays each having an incremental decrease in throughput capacity, but an increase in the depth and reliability of the data (Fig. 1a). The assays are grouped into four phases, each providing a new more detailed level of information about investigated compounds. The combination of all phases, including the optimization of the upstream steps based on downstream results (see feedback connections in Fig. 1a) was carried out to provide reliable characterization of the compound effect *in vivo*.

**Fig. 1:**
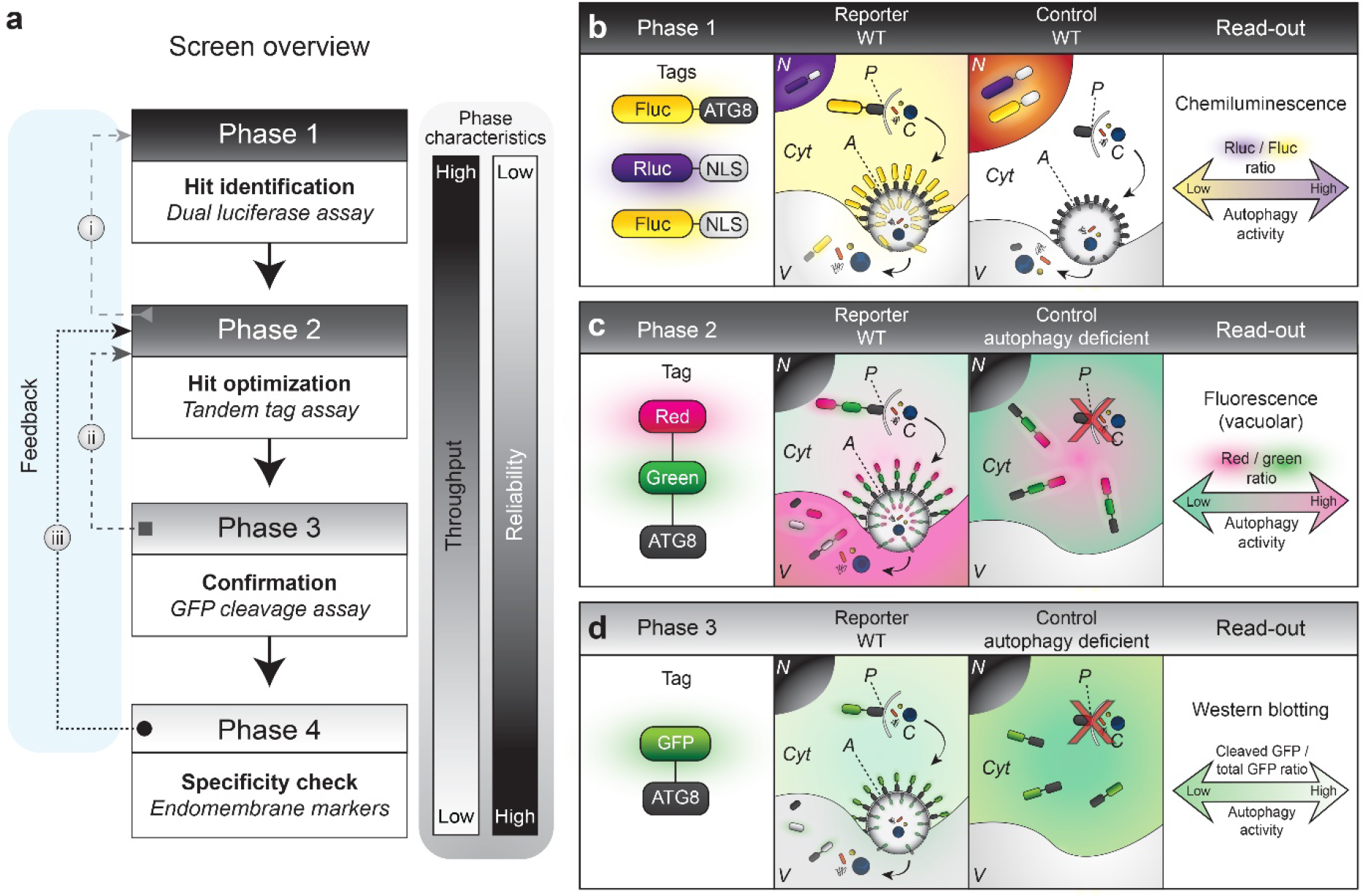
Multiphasic screen for autophagy-modulating compounds. **a**, The screen comprises four interconnected phases that allow identification of the best candidate compounds and verification of their specificity. The first phase is performed in a high-throughput mode, testing a library of compounds on cells expressing a reporter construct. The assay is performed in lysates of treated cells and can provide quantitative end-point read-outs. The assay of the second phase is performed *in planta* and has lower throughput as it employs advanced microscopy. It provides real-time quantitative read-out of autophagy activity and morphological data. While the read-out in the first two phases is based on delivery of autophagosomal membranes to the lytic vacuole, the third phase quantifies degradation of autophagic cargo, i.e. completion of the autophagic flux. Phase four is an essential step of the screen that verifies specificity of the compound’s effect to autophagy *in planta* using transgenic endomembrane marker lines. Importantly, results obtained in each phase are used for optimization of the previous steps: optimization of the concentration range for future screens (dashed arrow, i); estimation of efficacy and timing of cargo delivery versus its degradation (dashed arrow, ii); optimization the of concentration range of compounds for autophagy-specific effect (dashed arrow, iii). **b**, In the first phase a high throughput screen of compounds modulating autophagy is performed using the dual luciferase assay. **b**, reporter: wild-type (WT) tobacco BY-2 cells co-expressing firefly luciferase (Fluc) fused to ATG8 and *Renilla* luciferase (Rluc) fused to nuclear localization signal (NLS). Upon activation of autophagy, Fluc-ATG8 incorporates into autophagosome membranes and is delivered to the vacuole where Fluc activity is lost, while Rluc-NLS levels are unaffected. **b**, control: WT BY-2 cells co-expressing Fluc-NLS and Rluc-NLS, which are unaffected upon autophagy upregulation. The read-out for phase one is the ratio between chemiluminescence intensities of Rluc and Fluc substrates measured in cell lysates. **c**, In the second phase, delivery of ATG8 to the lytic vacuole is assessed using confocal microscopy. For this the tandem tag (TT) of red (TagRFP) and green (mWasabi) fluorescent proteins that are fused to ATG8 is expressed in a WT background (reporter) or in an autophagy-deficient background (control). Upon activation of autophagy, the TT-ATG8 is delivered to the vacuole in the reporter line, where green fluorescence is quenched by the acidic pH, while red fluorescence persists. The ratio between red and green fluorescence intensities in the vacuole serves as the read-out and is proportional to the rate of autophagosome delivery to the lytic compartment. **d**, In phase three the canonical GFP-cleavage western blot assay is implemented to assess completion of autophagic flux. For the assay, GFP-ATG8 is expressed in either a WT (reporter) or in autophagy-deficient background (control). Upon activation of autophagy in the reporter line, the GFP-ATG8 fusion is delivered to the vacuole and cleaved. Thus the ratio of cleaved *vs* total GFP provides the read-out for autophagy activity. Abbreviations, A = autophagosome; C = cargo; Cyt = cytoplasm; N = nucleus; P = phagophore; V = vacuole.

### Phase one – hit identification

The first phase was performed in the high-throughput mode using tobacco (*Nicotiana tabacum L.* cv Bright Yellow 2) cell cultures^16^ expressing chemiluminescent reporters for autophagic flux (Fig. 1b). Transgenic BY-2 cells were incubated and treated in 96-well format, lysed in the same plate and used for quantitative measurement of luminescence. In this assay the reporter line co-expressed *Renilla* luciferase (Rluc) fused to a nuclear localization signal (NLS) and firefly luciferase (Fluc) fused to ATG8. Upon activation of autophagy, Fluc-ATG8 was delivered to the lytic vacuole, leading to the loss of its activity and consequently resulting in an increase of the ratio of Rluc/Fluc. In the control line both luciferases are targeted to the nuclei, where they are protected from sequestration into autophagosomes. The dual luciferase assay permits measurement of both response (Fluc-ATG8 for the reporter line and Fluc-NLS for the control line) and reference (Rluc-NLS) luminescence in the same cells, thus removing possible bias originating from cell-to-cell fluctuations of expression or differences in cell number per sample. Additionally, absolute values of the Rluc luminescence can be used as a crude estimation of cell viability, as it generally dramatically decreases in lysates of dying cells.

The growth and treatment conditions were preliminarily optimized using known inducers and inhibitors of autophagy, e.g. AZD8055^17^, benzothiadiazole (BTH)^18^, and Concanamycin A (ConA)^19^ (Supplementary Fig. 1a, b and data not shown). Importantly, data from the viability assay of phase two (described below) provided a critical insight into an average range of compound concentrations that are not toxic for BY-2 cells (Fig. 2a). Therefore, it is advisable to perform a screen of a relatively small number of compounds first to estimate the optimal average concentration range for the library of interest (feedback connection i, Fig. 1a). As a proof of concept, a set of 364 small synthetic compounds of the Plasma Membrane Recycling compound set A (PMRA)^20^ and Plasma Membrane Recycling inhibitors in Pollen (PMRP)^20^ libraries was screened in this study and a total of 19 enhancers and 5 inhibitors of autophagy were chosen for further testing (Supplementary Fig. 1b).

**Fig. 2:**
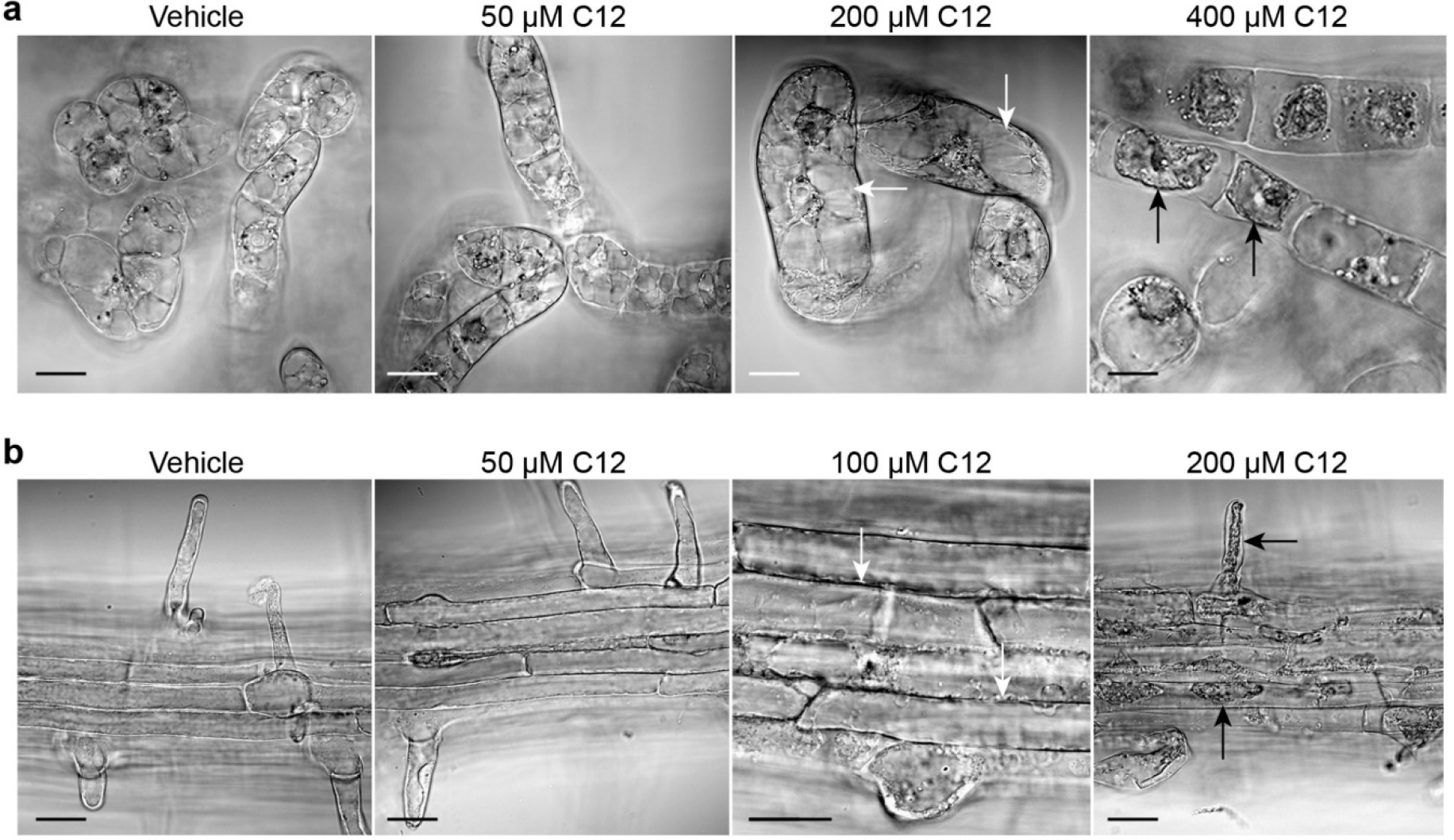
Phase two — viability testing. **a**, Tobacco BY-2 cell cultures and **b**, roots of 5-d-old Arabidopsis seedlings exposed for 4 h to a concentration gradient of compound 12 (C12) dissolved in 1% DMSO (vehicle). Viability is assessed at multiple timepoints using DIC light microscopy. Swollen (white arrows, **a**) and collapsed protoplasts (black arrows, **a**, in BY-2 cells at 200 and 400 µM concentrations, respectively. Arabidopsis roots with wrinkled (white arrows, **b**) or collapsed protoplasts (black arrows, **b**) at 100 and 200 µM concentrations, respectively. Scale bars, 20 µm (**a-b**).

### Phase two – hit optimization

To enhance the efficiency of the screen and promote an early elimination of compounds that do not reveal *in vivo* activity, *in planta* assays were performed early in the screen, already at the second phase.

#### Solubility assay

The first estimation of the applicable concentration range for the compounds selected in phase one was performed by assessing their solubility in the media used for growing cell cultures and seedlings under the corresponding growth conditions. For each selected compound we tested the concentration used in the first phase and additional concentrations that were several-fold greater. Additionally, at this stage it is advisable to examine the predicted *in vitro and in vivo* stability of the selected compounds under the treatment conditions^21^. This will allow correct handling of the compounds based on factors such as sensitivity to aqueous solutions, light or pH during upcoming experiments and omit compounds predicted to be unstable.

#### Viability assay

After estimating the range of possible concentrations of a compound, predicted by physical characteristics including but not limited to: molecular weight (MW), acid-base disassociation constant (p*K*a) and partition coefficient (logP)^22^, it is essential to identify the maximal concentration that is not cytotoxic, but still influences autophagy. Each compound at its maximal concentration selected based on the results of the solubility assay, along with serial dilutions were used for treatment of BY-2 cell cultures and Arabidopsis seedlings. Viability of the plant material was assessed by phenotyping specimens at 0–24 h of the treatment (depending upon the potency of each compound) using differential interference contrast (DIC) microscopy. Only the concentrations not causing any visible changes in cell morphology were considered for further assays.

An example presented in Fig. 2 illustrates the dose-dependent toxicity of the compound 12 (C12) after a 4 h treatment. At 50 µM, C12 had no observable effect on the BY-2 cells (Fig. 2a), whereas the 200 µM treatment caused swelling of the cells (white arrows, Fig. 2a) and 400 µM C12 led to protoplast collapse (black arrows, Fig. 2a). Interestingly, we observed that on average, root cells of Arabidopsis seedlings were considerably more sensitive to the compound treatments than BY-2 cells (Fig. 2). For instance, a 100 µM treatment of C12 resulted in wrinkled protoplasts in the epidermal root cells of Arabidopsis seedlings (white arrows, Fig. 2b), whereas 200 µM C12 led to protoplast collapse (black arrows, Fig. 2b). The difference in the sensitivity of the model system to the same compounds is an important factor to be assessed for phase one optimization (feedback arrow i, Fig. 1a).

#### Tandem tag assay

The first phase provides high-throughput quantitative data about delivery rate of the ATG8 fusion to the lytic compartment upon treatment of plant cell culture. Although this readout usually correlates well with autophagy activity, it still might provide false positive and false negative results. The Tandem Tag (TT) assay not only allows the assessment of time and dose-dependent delivery rate of the ATG8 fusion to the lytic compartment upon treatment with the compounds in *in planta*, but also yields important morphological data on autophagosome formation and confirms that the observed changes in ATG8 localization are autophagy-dependent. For this assay, ATG8 fused to the fluorescent tandem tag consisting of TagRFP and mWasabi^13^ was expressed in wild-type (WT) and autophagy-deficient Arabidopsis plants (reporter and control lines for autophagic flux detection, respectively; Fig. 1c). In the low pH of the vacuole, mWasabi green fluorescence is quenched more efficiently than TagRFP red fluorescence and therefore the ratio of the red to green fluorescence intensities in the vacuoles can be used as a read-out for autophagy activity, with higher ratios indicating increased autophagic flux.

Importantly, the basal autophagy level and compound sensitivity vary significantly depending on the cell type and differentiation stage. We observed the most reproducible results when performing microscopy experiments on epidermal root cells located at the beginning of the differentiation zone (DZ, Fig. 3a).

**Fig. 3:**
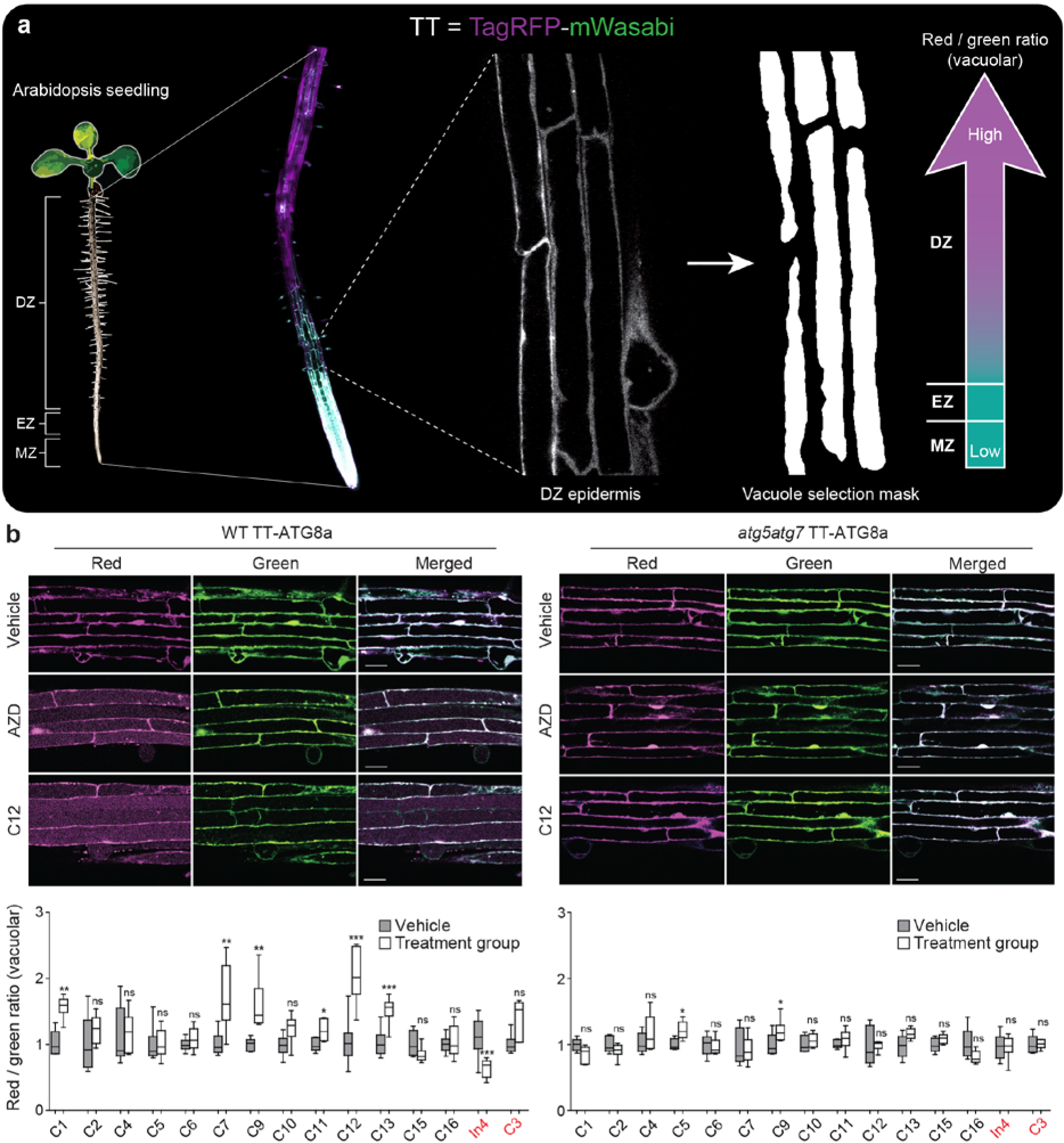
Phase two continued — hit verification and optimization *in planta*. **a**, Roots of Arabidopsis seedlings (5-d-old) expressing the Tandem Tag (TT; TagRFP-mWasabi) fused to ATG8a are utilized for hit optimization. The epidermal layer at the beginning of the root differentiation zone (DZ), located above the elongation and meristematic zones (EZ and MZ, respectively) is scanned via confocal microscopy. Images are then processed by our script to automatically quantify red (TagRFP) and green (mWasabi) fluorescence intensities in the vacuoles and perform statistical analysis of red to green ratios *vs* time or concentration. **b**, The TT-ATG8a vector is expressed in wildtype (WT) and *atg5atg7* Arabidopsis lines to determine the effects of each compound (C1-C16, In4) on autophagic flux. Treatments represented by red text (In4 and C3) are performed on seedlings exposed to nitrogen starvation for 24 h. Statistical information: n *≥* 6; *\*\*\* P <* 0.005*; ** P* < 0.01; *\* P <* 0.05; ns = non-significant (Student’s t-test); error bars represent SEM. Scale bars, 15 µm (**b**).

Firstly, we performed an estimation of the optimal concentration and treatment duration. For each compound, a range of concentrations below the cytotoxicity level identified in the viability assay was used to treat the TT reporter line only. The time-resolved changes in vacuolar fluorescence were imaged using confocal microscopy. The obtained data were originally quantified manually through the selection of small regions of the vacuoles using the DIC channel and then later automatically processed by our designated ImageJ-based macro and R script (Figs. 3, Supplementary Fig. 2a, Supplementary File 1). Next, the optimal treatment conditions for each compound were verified using both control (*atg5atg7* TT,) and reporter TT lines and the appropriate vehicle control (Fig. 3b). Figure 3b illustrates the effect of a set of compounds on red/green fluorescence, indicating increased autophagic flux. The assay revealed that under the tested conditions several compounds cause a significant change of the TT ratio compared to the vehicle treatment in the reporter line (WT TT-ATG8a, Fig. 3b), but not in the control line (*atg5atg7* TT-ATG8a, Fig. 3b).

The final optimization of the treatment conditions is carried out using the TT assay as the last step of phase four (feedback loop iii, Fig. 1a).

The TT assay in BY-2 cultures is less straightforward compared to Arabidopsis, due to the lack of corresponding autophagy-deficient control cell lines. Nevertheless, it still can be used by combining TagRFP-mWasabi-ATG8 and TagRFP-mWasabi-NLS as a reporter and control line for autophagic flux quantification, respectively (Supplementary Fig. 2b).

### Phase three – verification of autophagic flux completion

While the first two phases provide information about autophagy-dependent delivery of ATG8 to the lytic compartment, it remains unclear whether the final step of autophagic flux, i.e. cargo degradation, also takes place. The canonical assay to assess this is the GFP-ATG8 cleavage assay^23,24^.

Arabidopsis plants expressing GFP-ATG8a in WT or autophagy-deficient backgrounds are used as a reporter and control lines, respectively, for autophagic flux quantification (Fig. 1d). Upon induction of autophagy, GFP-ATG8 is delivered to the vacuole at a higher rate, where it is first cleaved by lytic proteases, leading to transient accumulation of the cleavage product GFP. Western blot analysis of protein extracts allows the estimation of a ratio between total and cleaved GFP in plant cells, thus revealing the relative level of autophagic flux (Fig. 1d).

Importantly, tissue and organ-specific differences in the levels of 2×35S-driven expression of GFP-ATG8 and/or basal autophagic flux can considerably skew results. For example, protein extracts from whole seedlings compared to the root differentiation zone of the same lines under the same treatment conditions reveal significant differences in the read-out (Fig. 4a). We established a designated protocol for the GFP-ATG8 cleavage assay for the Arabidopsis root differentiation zone and obtained reproducible results consistent with the results of the TT assay (Fig 4b, c; Fig 3b). Notably, the combination of GFP-ATG8 cleavage assay results with morphological and quantitative data obtained via the TT assay may indicate which step of the autophagy pathway is affected by the investigated compound (feedback arrow ii, Fig. 1a). For example, inhibitor 4 (In4) had an accumulation of cytoplasmic TT-positive puncta at high concentrations, suggesting fusion of the autophagosomes to the vacuole and autophagic flux was disrupted (data not shown). Interestingly, slightly greater concentrations of putative autophagy modulators or longer incubation times were needed for efficient GFP-ATG8 cleavage assay, compared to the TT assay. This might be caused by differences in the cells types that were included in each assay (only epidermal cells for the TT assay *vs* all cell types for the GFP-ATG8 cleavage assay) or differences in the sensitivity of the methods. Alternatively, it might indicate that GFP-ATG8 fusion hydrolysis does not occur immediately after delivery of the autophagosomes to the lytic compartment.

**Fig. 4:**
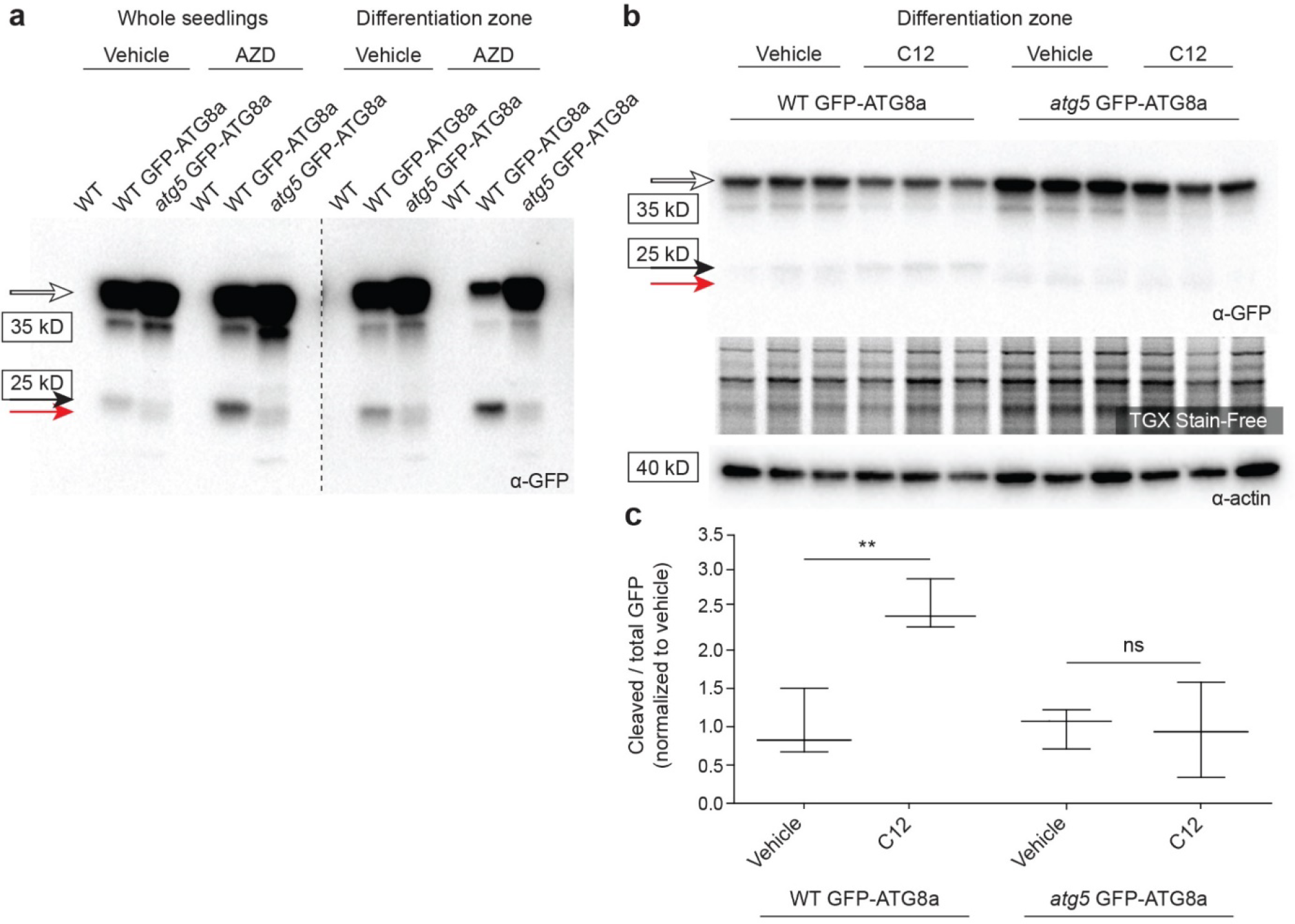
Phase three — validation of autophagic flux completion. **a**, Green fluorescent protein (GFP) cleavage assay for Arabidopsis whole seedling and root differentiation zone protein extracts from the following lines: wild-type (WT), GFP-ATG8a expressed in WT (WT GFP-ATG8) and *atg5* KO (*atg5* GFP-ATG8a). Treatments were applied for 6 h prior to protein extraction and included a vehicle control (1% DMSO) and 0.5 µM AZD8055 (AZD). **b**, Immunoblotting with α-GFP of differentiation zone protein extracts from WT GFP-ATG8a and *atg5* GFP-ATG8a lines (three independent replicates per treatment and genetic background) treated with 1% DMSO (vehicle) or 30 µM of compound 12 (C12) for 4h. Bio-Rad TGX Stain-Free gels and α-actin are utilized as loading controls. **a, b**, GFP-ATG8a fusion band (white arrow); Cleaved GFP (black arrow); lower molecular weight band (red arrow). Dashed line indicates membrane splicing. **c**, Cleaved versus total GFP ratios comparing vehicle and 30 µM C12 treatment in WT GFP ATG8a and *atg5* GFP-ATG8a lines. Statistical information: n *=* 3; ** *P* < 0.01; ns = non-significant (Student’s t-test); error bars represent SEM.

### Phase four – specificity check

Autophagy is integrated tightly with endo- and exocytic pathways and the trafficking routes to the vacuole^25–27^. Thus, it is probable that putative modulators of autophagy identified in the previous phases of the screen influence the endomembrane system and thus activate or inhibit autophagy indirectly and have significant off-target effects, as was recently shown for some characterized compounds^10,11^.

To minimize this possibility, we designed the fourth phase of the screen where the effect of each candidate compound on endomembrane compartments is tested *in planta* using a library of fluorescent marker lines (Supplementary Table 3). The library comprises the minimal set representing the key components of the endomembrane system including: endoplasmic reticulum (ER), Golgi apparatus (GA), *trans*-Golgi network (TGN), multi-vesicular bodies (MVBs), vacuole, mitochondria, peroxisomes, plasma membrane, and endo- and exocytic pathways.

Each compound was tested on the set of the marker lines using the conditions optimized in the second and third phases. If detectable morphological changes were observed for any of the markers, a range of lower compound concentrations and treatment times were tested. This allowed us to estimate the maximum possible concentration at which a compound did not change the morphology of any endomembrane marker. The identified concentration was then tested using the TT assay to assess its impact on autophagy activity (feedback arrow iii, Fig. 1a).

Figure 5 provides an example of optimization for C12 at conditions originally optimized in phases two. A gradient of C12 concentrations was tested using an ER marker line (Fig. 5a). Concentrations of C12 at 60 µM and above caused significant ER stress, resulting in a disrupted cisternae network and formation of swollen GFP-positive bodies (white arrows, Fig. 5a; Supplementary Video 1). C12 at 30 µM or less did not alter ER morphology (Fig. 5a). Thus, C12 concentrations ranging from 7.5 µM to 30 µM were tested using the TT and GFP-ATG8a cleavage assay and showed a significant impact on autophagic flux (Fig. 5b, c).

**Fig. 5:**
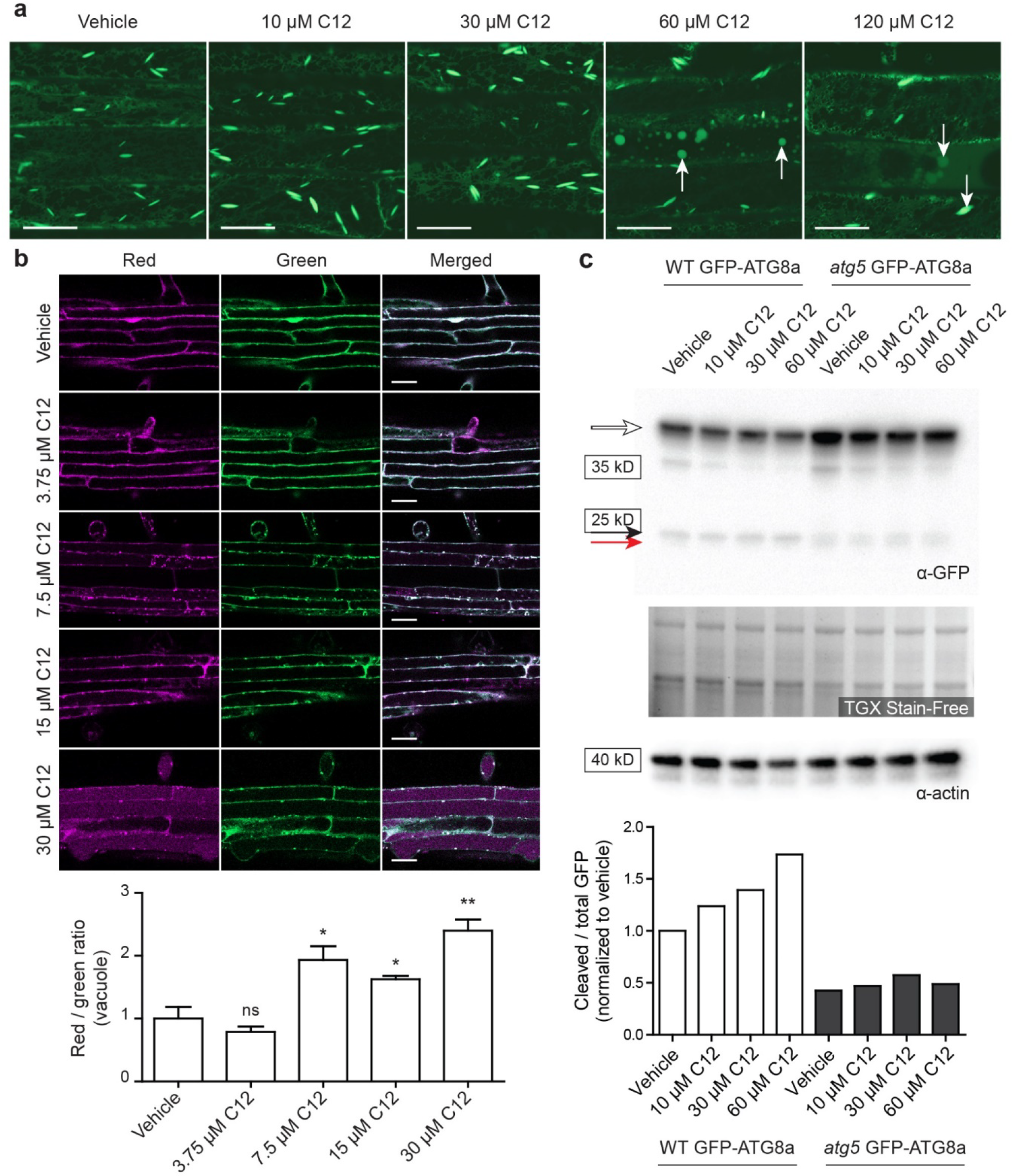
Phase four – identification of autophagy-specific compounds and conditions. **a**, HDEL-GFP Arabidopsis seedlings exposed to a gradient of compound 12 (C12) concentrations to assess its phenotypic effects on the endoplasmic reticulum (ER). Swollen ER bodies (white arrows) and disrupted ER networks occur at higher concentrations (60 and 120 µM). **b**, Tandem tag (TagRFP-mWasabi)-ATG8a reporter lines treated with a range of C12 concentrations. **c**, The GFP-cleavage assay for a gradient of C12 treatments. Western blot detection of GFP in protein extracts prepared from differentiation zone of WT GFP-ATG8a and *atg5* GFP-ATG8a lines. Bio-Rad TGX Stain-Free gels and α-actin immunoprobing are used as loading controls. GFP-ATG8a fusion band (white arrow); cleaved GFP (black arrow); lower molecular weight band (red arrow). Scale bars, 15 µm (**a**), 20 µm (**b**). Statistical information (**b**): n = 3; ** *P* < 0.01; * *P* < 0.05; ns = non-significant (Student’s t-test); error bars represent SEM.

## Discussion

We describe a novel robust pipeline designed for the identification of chemicals specifically modulating plant autophagy. Our approach resulted in hits with minimal off target effect and provides data essential for further characterization of bioactive candidates. One of the most important features of the new pipeline is the combination of a high-throughput screen utilizing cell cultures with further verification steps performed *in planta*. This approach simultaneously enables screening large numbers of compounds and eliminates false positive candidates that are active in cell cultures, but do not have activity *in planta* due to differences in cell permeability, stability in media used for cell culture and plant growth or cell-type-specific responses. Furthermore, combining the use of *Nicotiana tabacum* cell cultures and Arabidopsis seedlings provides a first estimation of potential conservation of targets in plant species belonging to different families. Interestingly, to our experience, the average concentration range of compounds applicable for BY-2 cells was significantly higher than the concentration range suitable for Arabidopsis plants, indicating differences in responses between the two models. It is also possible however, that BY-2 cells would respond similarly at the same concentrations with increased exposure times, but more detailed kinetic experiments would be required to test this hypothesis. Nevertheless, the difference observed is a useful starting point for future screens utilizing either of the model systems. Notably, the absolute values of the applicable concentration ranges must be individually adjusted to a specific chemical library, as the activity of compounds might vary due to purity, age, structural stability and storage differences.

As further characterization of identified autophagy modulators is costly and labor-intensive^28,29^, it is important to identify bioactive chemicals that specifically modulate autophagy under physiological conditions. To achieve this, we designed a multi-faceted approach in which the first three phases involved advanced fluorescent microscopy, biochemistry and cell and molecular biology methods. This approach provides quantitative and morphological data on autophagy pathway activity and is optimized to assess the effect of a chemical on several key aspects of the pathway. This characterization not only ensures reliability of the conclusions about effects on the plant autophagy, but also provides an indication of the affected stage of the autophagy pathway, e.g. phagophore formation, cargo delivery or cargo degradation. While assays of the first three phases are focused on assessing the chemicals’ effect on autophagy activity, the fourth phase puts the activity into a broader context. Due to the overwhelming complexity of the network, it is frequently overlooked that autophagy is an integral part of the endomembrane trafficking system^27,30,31^. The assays in the fourth phase identify treatment conditions under which candidate compounds significantly influence autophagy activity without causing detectable aberrations in other components of the trafficking machinery. Although informative, these assays do not provide a definite proof of autophagy specificity of the compound. A more detailed estimation of potential off-targets of the compound can be achieved via assessment of its effect on loss-of-function mutants of the predicted target and predicted off-targets^32^.

The pipeline described here is highly efficient in identifying specific regulators of plant autophagy and optimizing the treatment conditions for plants grown in laboratory conditions, thus providing a powerful tool for fundamental plant autophagy research. Further characterization of the identified compounds can include conventional methods for target identification^28,33^ and/or direct testing of compounds on plants grown in soil. The first approach will provide an in-depth mechanistic understanding of the modulator’s activity – information valuable for optimization of specificity and potency, as well as predicting its applicability for other species, including animal and human disease models. The second approach might potentially streamline direct application of the compounds in the field, enabling the use of chemical transient modulators of plant autophagy as potent regulators of crop fitness^1,4,34^, thus benefiting modern agriculture.

## Methods

### Genetic constructs

Maps and sequences of all constructs used in this study can be found in Supplementary Table 1. Information about the primers used for cloning is available in Supplementary Table 2.

To generate the entry clone AM425, AtATG8a CDS was amplified from total cDNA of Arabidopsis seedlings using primers AM18/AM19. The obtained attB-flanked amplicon was then recombined with the Gateway^TM^ pDONR^TM^/Zeo vector (ThermoFisher) using Gateway^TM^ BP Clonase II enzyme mix (ThermoFisher).

Firefly luciferase (Fluc) and *Renilla* luciferase (Rluc) coding sequences were amplified using the primer pairs PB31/PB32 and PB33/PB34 and vectors pRD400 35S::Fluc and pRD400 35S::Rluc^35^, respectively. The amplicons were inserted using KpnI/AscI restriction digestion sites into a pMDC32 vector^36^ that was previously modified to replace hyg resistance gene with nptII sequence. The resulting binary Gateway^TM^–based destination vectors were named AM616 and AM618 (Supplementary Table 1).

AM439 pMDC kan Fluc AtATG8a was obtained by recombining the entry clone AM425 (see above) with AM616 pMDC kan Fluc using Gateway^TM^ LR Clonase II enzyme mix (ThermoFisher).

The synthetic sequence of attB-flanked SV40 NLS (Eurofins Scientific, Sweden) was recombined with the Gateway^TM^ pDONRTM/Zeo vector (ThermoFisher) using Gateway^TM^ BP Clonase II enzyme mix (ThermoFisher), producing entry clone CC3.

The CC3 entry clone was then recombined with the AM618 vector to produce an intermediate clone containing the sequence of 2×35S::*Renilla*-NLS fusion::NosT. This sequence was amplified using the 5’-phosphorylated primers CC1 and CC2. To generate the CC8 destination clone, the amplicon was then blunt-end ligated into the AM439 destination clone linearized with the PmeI restriction digestion enzyme (NEB).

The CC10 clone was obtained by introducing two stop codons between the Fluc and AtATG8a utilizing The QuikChange^®^ site-directed mutagenesis (Agilent) and primers CC7/CC8.

The sequence of TagRFP-mWasabi was amplified from the plasmid kindly provided by Zhou and colleagues^13^ using the primers PB15/PB16. The amplicon was inserted into pMDC32 or pMDC kan to produce destination binary vectors, AM613 and AM614, respectively. These two vectors were then recombined with the entry clone AM425 using Gateway^TM^ LR Clonase II enzyme mix (ThermoFisher) to produce AM434 and AM435 clones, respectively.

### Plant material

The description of stable transgenic lines used in this study is available in Table S3.

### Plant growth and transformation

Arabidopsis seeds were surface sterilized using a bleach solution comprising 10% Klorin (Sweden) with 0.05% Tween-20 for at least 20 min, washed three times with sterile MQ water and plated on MS plates: 0.5xMS including vitamins (M0222, Duchefa), supplemented with 10 mM MES (M1503, Duchefa), 1% sucrose, 0.8 % Plant agar (P1001, Duchefa), pH 5.8. Seeds on plates were then vernalized in darkness at 4°C for 24-48 h. After vernalization, plates were incubated vertically under long day growth conditions (16 h of 150 µM light at 22°C, 8 h dark at 20°C).

For growth in soil (2001, Hasselfors Garden), seeds were surface sterilized as described above, vernalized in MQ water in Eppendorf tubes and then transferred directly into soil. Plants were grown under the long day conditions as described above.

For transformation, Arabidopsis plants were grown in soil for approximately 5 weeks, the first inflorescence was clipped to stimulate production of auxiliary inflorescences. All genetic constructs used for transformation were first introduced into *Agrobacterium tumefaciens* strain GV3101^37^. The floral dip transformation was performed as previously described^38^. Transformants were selected on the 0.5x MS medium as described above supplemented with 400 µg/ml timentin (T0190, Duchefa), 250 µg/ml cefatoxime (C0111, Duchefa) and either 35 µg/ml hygromycin B (H0192, Duchefa) or 50 µg/ml kanamycin (K0126, Duchefa). Lines showing a 3:1 segregation pattern in the T2 generation were used for establishing homozygous T3 lines.

For testing potential inhibitors of autophagy, nitrogen starvation was induced in seedlings by incubating them in liquid nitrogen-free MS medium: 0.5x MS basal salt micronutrient solution (M0529, Sigma), 1% sucrose, 3 mM CaCl2, 1.5 mM MgSO4, 1,25 mM KH2PO4, 5 mM KCl, 0.8% Plant agar, pH 5.8. Seeds were germinated on standard 0.5x MS medium as described above. Seedlings were grown on vertically positioned plates under long day conditions for 4– 5 d before being transferred into wells of 6-well tissue culture plates with 3 ml of nitrogen-free MS medium. The medium was gently pipetted on top of the roots to ensure their complete submersion. Nitrogen starvation effect was clearly detectable after 18 h of treatment.

### BY-2 cell culture growth and transformation

Tobacco BY-2 suspensions were subcultured under sterile conditions every 7–10 d using the standard medium: 1x MS including vitamins (M0222, Duchefa), 0.02% KH2PO4, 0.2 mg/L 2,4-D, 3% sucrose, pH 5.6. The cultures were incubated in Erlenmeyer flasks at 22°C in the darkness, on a gyratory shaker at 120 rpm.

For transformation of BY-2 cells, overnight cultures of transgenic *Agrobacterium tumefaciens* GV3101 were grown overnight at 28°C, 180 rpm, in 3 ml of LB medium supplemented with 50 µg/ml rifampicin and 50 µg/ml kanamycin. Bacterial cells from the overnight cultures were spun down for 15 min at 4 000 g, resuspended in 2 ml of BY-2 medium supplemented with 150 µM acetosyringone (D134406, Sigma-Aldrich) and incubated at 28°C, 120 rpm for one hour. The bacterial cells were then added to 4 ml of 4-d-old BY-2 culture to the final OD_600_ of *Agrobacterium* = 0.375. The mixture was transferred on plates with BY-2 medium solidified with 0.8% agar. The plates were kept at 28°C for 48 h, after which the cells were gently transferred on plates containing 400 µg/ml timentin (T0190, Duchefa), 250 µg/ml cefatoxime (C0111, Duchefa) and either 40 µg/ml hygromycin B (H0192, Duchefa) or 50 µg/ml kanamycin (K0126, Duchefa). The first *calli* were typically visible after 2 weeks of growth on the selection plates.

### Treatment of Arabidopsis seedlings for microscopy

For microscopy experiments, treatment of Arabidopsis seedlings with candidate compounds was performed in 6-well tissue culture plates. Five-d-old seedlings grown on vertical plates were transferred into wells containing 3 ml of standard or nitrogen-free 0.5x MS medium supplemented with a compound or the corresponding concentration of the vehicle (DMSO). The seedlings were then submerged by gently pipetting the medium on the roots. The plates were sealed with a strip of Saran wrap and incubated in the growth cabinet under long day conditions for the designated time. For microscopy, the seedlings were mounted in the corresponding treatment medium.

Prior to the viability assay, solubility was assessed in small volumes of MS medium to exclude concentrations at which the compound precipitated. Viability assays were performed using Zeiss AxioImager 2 epifluorescent microscope equipped with DIC (Nomarski) optics. Images were acquired using ZEN blue software (version 2.1, Carl Zeiss).

### Treatment of BY-2 cells

For the primary screen, 3-d-old BY-2 cell culture was pipetted into 96-well plates (140 µl/well), using wide orifice tips. The plates were kept on an orbital shaker (4 mm) at 640 rpm, at 25°C for 24 h to let cells recover from the mechanical stress. One µl of each compound taken from the PMRP^39^ and PMRA^20^ libraries was pipetted into the plates with the BY-2 cells. One µM AZD8055 and one mM BTH were used as positive controls for upregulation of autophagy, 1% DMSO was used as a negative control. The cells were treated while shaking at 25°C, in the dark for 24 h, after which 50 µL of 4x Passive Lysis Buffer (Dual Luciferase Assay kit, E1910, Promega) was added to all wells and the plates were sealed and frozen at −80°C to rupture the cell walls. To measure the luciferase intensity values, the plates were thawed at RT, vortexed for 2 min and spun down at 4 000g for 5 min. Ten µl of each cell lysate were used for dual luciferase assay by applying 50 µL of freshly prepared LARII and Stop&Glo reagents of the Dual Luciferase Assay kit (E1910, Promega). The luciferase intensity values of firefly and *Renilla* luciferases were measured sequentially by collecting one second of integrated chemiluminescence intensity (using gain 240) after a delay of two seconds upon injection of each substrate (LARII and Stop&Glo respectively), using the Synergy^TM^ H4 Multi-Mode Microplate Reader (BioTek).

The obtained data was then analysed using R (version 3.5.2)^40^. The compounds were ranked according to their *Renilla* to firefly luciferase intensity ratio and to their relative luminescence unit (RLU) values for Rluc. The latter were used to infer the viability of the cells and low values (below 150000 RLU) were considered to correspond to cytotoxic effects from the treatment. Rluc RLU values produced by dying cell cultures will vary depending on detector sensitivity and therefore should be estimated prior to screening. Compounds inducing more than 2.5 fold increase in *Renilla* to firefly luciferase ratio compared to the DMSO control and Rluc RLU values above 150000, were selected as putative enhancers of autophagy. Compounds inducing a two-fold reduction of the *Renilla* to firely luciferase ratio and Rluc values above 150000 RLU were selected as putative inhibitors.

Treatment of BY-2 cells for further analyses was performed in 6-or 24-well plate format in 1– 3.5 ml of 4-d-old BY-2 culture. After transfer into the plate and before the start of a treatment, the cells were left to recover for 4–18 h. During treatment, the plates were kept at 25°C, 120 rpm in the dark. For sampling, the cells from each well were harvested onto a 50 µm nylon mesh, frozen and ground in liquid nitrogen and resuspended in 1x passive lysis buffer (Dual Luciferase Assay kit, E1910, Promega), using 2 volumes/mg of material. Ten µl of each cell lysate was then transferred into another plate and used for dual Luciferase assay applying a quarter of the recommended volume of each reagent of the Dual Luciferase Assay kit (E1910, Promega). The substrates were added by an automated pump of the FLUOstar Omega Microplate Reader (BMG LABTECH) directly prior to double orbital shaking at 100 rpm for 1 s and then a 2 s reading interval for each well. The gain adjustment was estimated in a pilot experiment using lysates of BY-2 cells treated with 1% DMSO, 500 nM AZD and 1 µM ConA.

### Tandem tag assay

The detailed protocol for TT assay is described in Supplementary File 1.

### Western blot

For the western blot analysis, Arabidopsis seedlings were grown on vertical plates as described above. Briefly, seeds were sawn in two rows per plate and the seedlings were grown for 5-7 d until the root became 4-5 cm long. For treatment, the plates were placed horizontally and flooded with liquid 0.5x MS medium supplemented with the corresponding compound. Treatments were performed in the growth cabinet under long day conditions. Upon the completion of the treatment, the excess liquid was poured out of the plates and the whole seedlings were harvested and ground in liquid nitrogen. Alternatively, the differentiation zone of the root was excised from approximately 50 seedlings using a scalpel blade, gently collected with the tweezers and briefly dried on a paper tissue to be then placed into a 1.5 ml Eppendorf and ground in liquid nitrogen with a plastic pestle. The powdered material was mixed 1:2 (w/v) with 2x Laemmli buffer^41^, the mixture was thoroughly vortexed and boiled for 5 min at 95°C. Samples were spun down at 13 000 g for 10 min, and 1–7.5 µl of the supernatant were loaded on precast gels (Bio-Rad). Proteins were transferred onto a PVDF membrane and blotted using 1:2000 a-GFP (Cat 632381, Clontech), 1:2000 a-actin (AS13 2640, Agrisera), 1:25 000 a-rabbit-HRP (AS09602, Agrisera), 1:8000 a-mouse-HRP (NA931, Amersham). The blots were developed using ECL Prime (Amersham) and GelDoc^TM^ XR (Bio-Rad). The intensities of the bands were quantified using Image Studio Lite (version 5.2, LI-COR) software.

### Statistical analysis and image preparation

Statistical analysis was performed using GraphPad Prism (version 7.03, GraphPad Software) unless otherwise stated. Figures were prepared using Adobe Photoshop and Illustrator CC (Adobe Inc.). Adjustments to brightness and contrast were applied equally to corresponding images within figures.

## Acknowledgements

This project was supported by grants from Olle Engkvist Foundation (to PVB, EAM and SR), Carl Tryggers Foundation (to EAM), MSCA IF (to EAM), the Swedish Foundation for Strategic Research (to PVB), the Swedish Research Councils VR and Formas (to PVB), the Knut and Alice Wallenberg Foundation (to PVB) and by the research programme “Trees and Crops for the Future” at the Swedish University of Agricultural Sciences. Postdoctoral fellowship of AND was provided by the Natural Sciences and Engineering Research Council (NSERC) of Canada. GRH is grateful to August T. Larsson Guest Researcher Programme for supporting his visits to the Swedish University of Agricultural Sciences.

## Author Contributions

AND performed most of the experiments for phases two through four and wrote a part of the manuscript. CC established and performed the primary screen in the phase one and established transgenic lines. KD helped establish the transgenic lines for the study. JAO contributed to designing the scripts for the automated vacuolar fluorescence intensity measurement. SBF contributed to the primary screen in the phase one. SR contributed to conceiving the project idea and the primary screen in the phase one. GRH assisted with phase four experiments. PVB secured funding and contributed to conceiving the project idea. EAM contributed to conceiving the project idea, supervised the project, performed some experiments and wrote a part of the manuscript.

## Materials & Correspondence

For material request and correspondence please contact Alyona Minina at

## Data availability

The data relevant to this article will be made available upon a personal request. Please note that some of the information about identified modulators of plant autophagy will be made public in upcoming publications.

## Supplementary information

**Supplementary Fig. 1:**
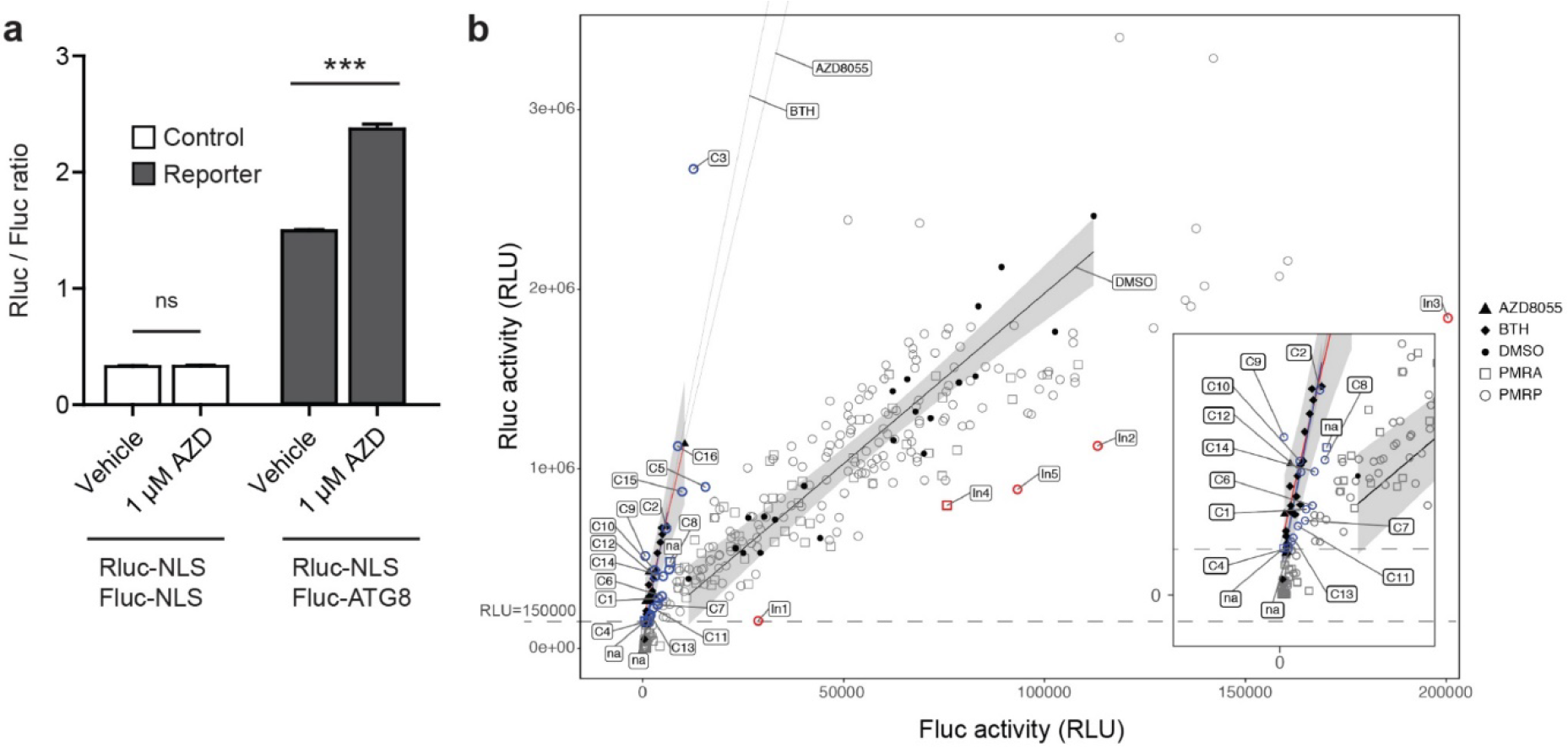
Dual luciferase reporter verification and hit identification in tobacco BY-2 cells. **a**, Dual luciferase assay validation. Control cell line was co-expressing *Renilla* luciferase (Rluc) and firefly luciferase (Fluc) fused to Nuclear Localization Signal (NLS), whereas the reporter line was co-expressing Rluc-NLS and Fluc fused to ATG8. Upon activation of autophagy by AZD8055 (AZD), the ratio of Rluc/Fluc increases in the reporter but not in the control line. Statistical information: n = 3; *** *P* < 0.0001; ns = non-significant (Student’s t-test); error bars represent SEM. **b**, Hit identification using the BY-2 reporter line. A screen of 364 compounds from the Plasma Membrane Recycling compound set A (PMRA) and Plasma Membrane Recycling inhibitors in Pollen (PMRP) libraries reveals several putative inhibitors (> 2-fold decrease in Rluc/Fluc ratio, highlighted in red) and enhancers (> 3.3-fold increase in Rluc/Fluc ratio, highlighted in blue). Enhancers with relative luminescence unit (RLU) values below 150000 were considered potentially cytotoxic and were excluded from further characterization. Lines and shaded areas represent the linear regressions and 95% confidence intervals for each of the control treatments, including DMSO (black), BTH (blue, inset) and AZD8055 (red, inset). Inset shows magnified part of the major chart. Please note that BTH and AZD8055 lines were extended and labelled for clarification.

**Supplementary Fig. 2:**
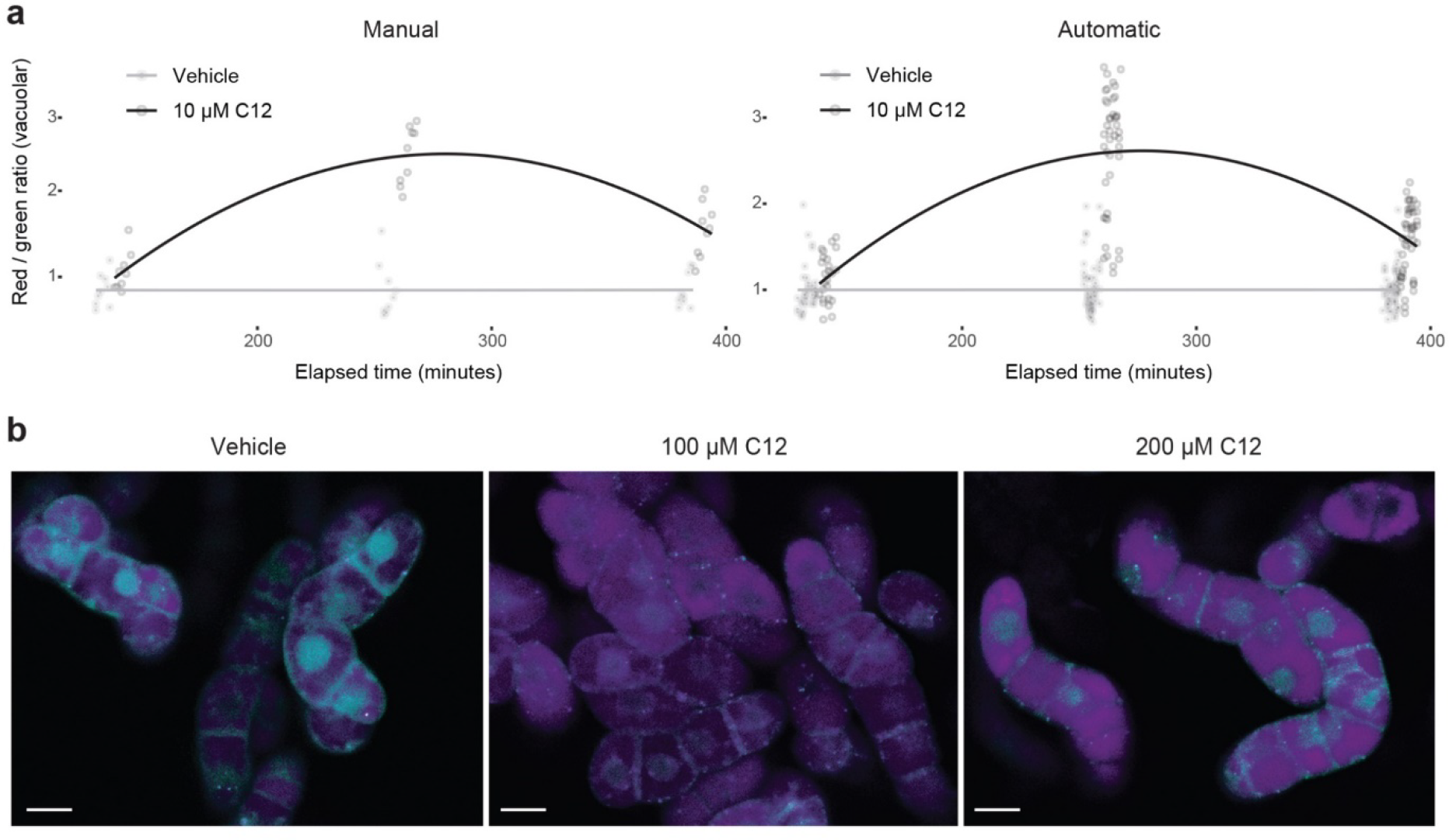
Tandem tag assay. **a**, Comparison of manual and automatic quantification of vacuolar red to green fluorescence intensities ratios in Arabidopsis seedlings expressing tandem tag (TagRFP-mWasabi) fused to ATG8a and treated with 10 µM compound 12 (C12) or vehicle (1% DMSO). **b**, An example of tandem tag detection in tobacco BY-2 cell cultures treated with the vehicle (1% DMSO), 100 µM C12 and 200 µM C12 for 4 h. Scale bars, 20 µM (**b**).

**Supplementary Video 1: Phase four examples.**

**a**, Timeseries and Z-stacks (**b**) of Arabidopsis seedling expressing the ER-marker HDEL-GFP. (**a**, **b**) Root differentiation zone epidermal cells from vehicle (1% DMSO), 10 µM compound 12 (C12) and 60 µM C12 treatment groups. Scale bars, 15 µm. Timeseries acquisition time: 2 minutes. Click on individual videos to start and stop playback.

**Supplementary Table 1.**
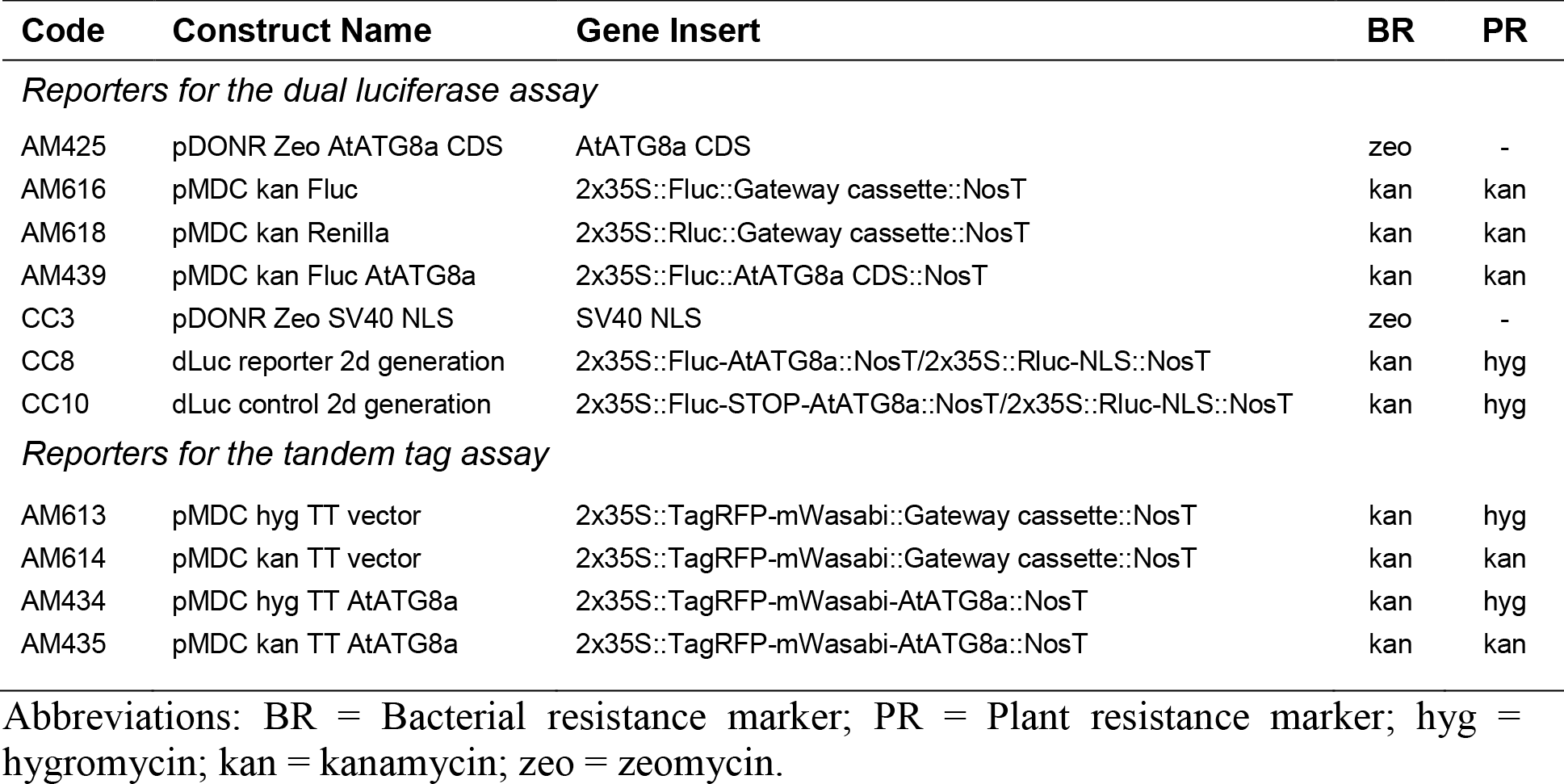
Constructs used in this study.

| Code | Construct Name | Gene Insert | BR | PR |
| --- | --- | --- | --- | --- |
| *Reporters for the dual luciferase assay* |  |  |  |  |
| AM425 | pDONR Zeo AtATG8a CDS | AtATG8a CDS | zeo | - |
| AM616 | pMDC kan Fluc | 2x35S::Fluc::Gateway cassette::NosT | kan | kan |
| AM618 | pMDC kan Renilla | 2x35S::Rluc::Gateway cassette::NosT | kan | kan |
| AM439 | pMDC kan Fluc AtATG8a | 2x35S::Fluc::AtATG8a CDS::NosT | kan | kan |
| CC3 | pDONR Zeo SV40 NLS | SV40 NLS | zeo | - |
| CC8 | dLuc reporter 2d generation | 2x35S::Fluc-AtATG8a::NosT/2x35S::Rluc-NLS::NosT | kan | hyg |
| CC10 | dLuc control 2d generation | 2x35S::Fluc-STOP-AtATG8a::NosT/2x35S::Rluc-NLS::NosT | kan | hyg |
| *Reporters for the tandem tag assay* |  |  |  |  |
| AM613 | pMDC hyg TT vector | 2x35S::TagRFP-mWasabi::Gateway cassette::NosT | kan | hyg |
| AM614 | pMDC kan TT vector | 2x35S::TagRFP-mWasabi::Gateway cassette::NosT | kan | kan |
| AM434 | pMDC hyg TT AtATG8a | 2x35S::TagRFP-mWasabi-AtATG8a::NosT | kan | hyg |
| AM435 | pMDC kan TT AtATG8a | 2x35S::TagRFP-mWasabi-AtATG8a::NosT | kan | kan |

Abbreviations: BR = Bacterial resistance marker; PR = Plant resistance marker; hyg = hygromycin; kan = kanamycin; zeo = zeomycin.

**Supplementary Table 2.**
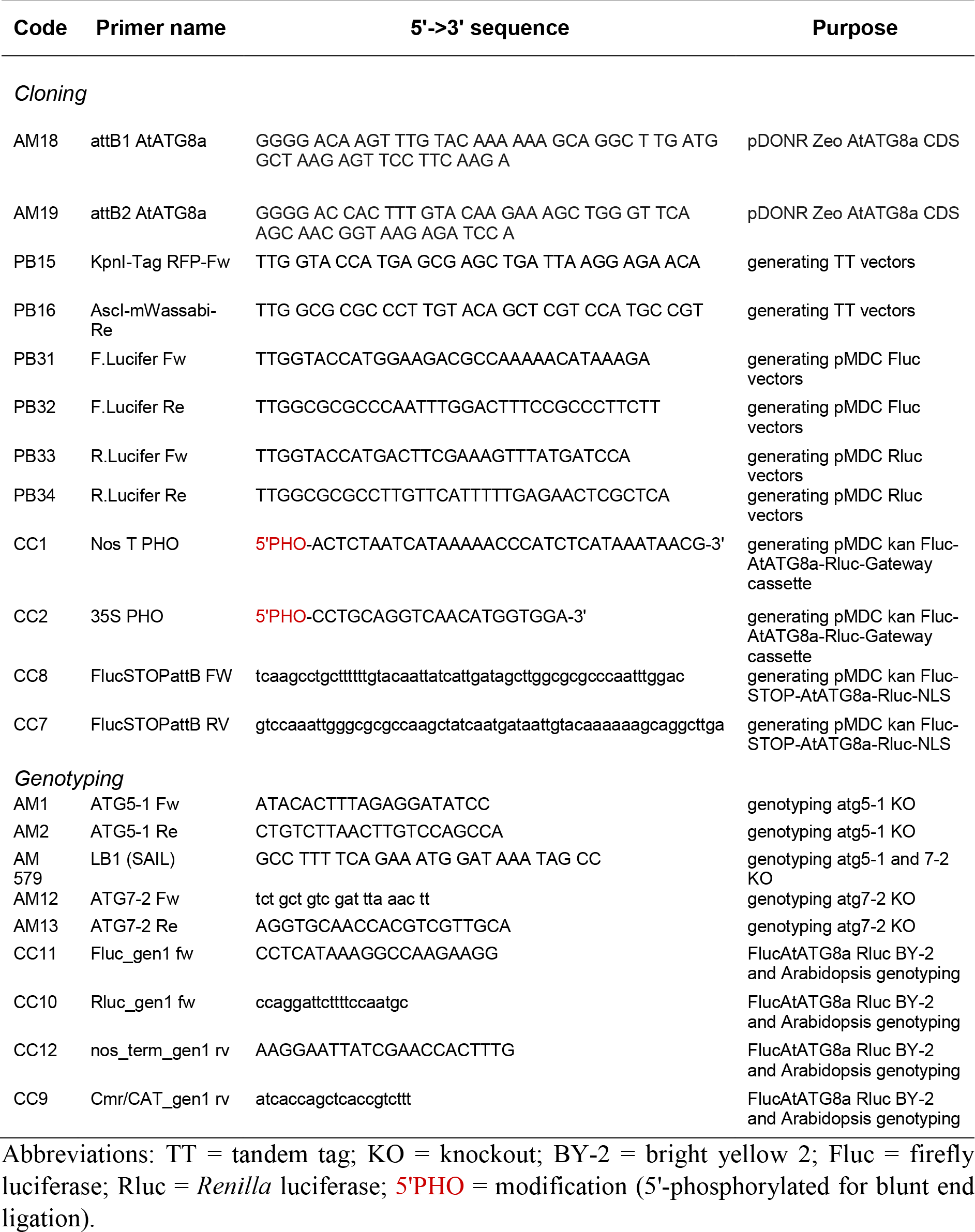
Primers used in this study.

**Supplementary Table 3.** Stable transgenic lines used in this study.

| Line | Transgene | BG | Acc | Gen | HO | PR | Origin |
| --- | --- | --- | --- | --- | --- | --- | --- |
| *Tobacco BY-2 Cell Lines* |  |  |  |  |  |  |  |
| dLuc reporter | 2x35S::Fluc-AtATG8a::NosT/2x35S::Rluc-NLS::NosT | WT | N/A | N/A | N/A | hyg | This study |
| dLuc control | 2x35S::Fluc-STOP-AtATG8a::NosT/2x35S::Rluc-NLS::NosT | WT | N/A | N/A | N/A | hyg | This study |
| TT reporter | 2x35S::TagRFP-mWasabi-AtATG8a::NosT | WT | N/A | N/A | N/A | kan | This study |
| TT control | 2x35S::TagRFP-mWasabi::NosT | WT | N/A | N/A | N/A | kan | This study |
| *Arabidopsis thaliana lines* |  |  |  |  |  |  |  |
| TT reporter | 2x35S::TagRFP-mWasabi-AtATG8a::NosT | WT | Col-0 | T3 | HO | hyg | This study |
| TT control | 2x35S::TagRFP-mWasabi-AtATG8a::NosT | atg5-1/atg7-2 | Col-0 | T3 | HO | hyg | This study |
| GFP-ATG8a reporter | 2x35S::GFP-AtATG8a | WT | Col-0 | N/A | HO | basta | (Thompson et al.)<sup>42</sup> |
| GFP-ATG8a control | 2x35S::GFP-AtATG8a | atg5-1 | Col-0 | F5 | HO | basta | (Minina et al.)<sup>34</sup> |
| 35S GFP-HDEL | 2x35S::GFP-HDEL | WT | Col-0 | N/A | HO | kan | (Nelson & Nebenfuhr)<sup>43</sup> |
| 35S MT-YFP | 2x35S::MT-YFP | WT | Col-0 | N/A | HO | kan | (Nelson & Nebenfuhr)<sup>43</sup> |
| 35S::NAG1-eGFP | 35S::NAG1-eGFP | WT | Col-0 | T3 | HO | N/A | (Grebe et al.)<sup>44</sup> |
| VHAa1::VHAa1-GFP | VHAa1::VHAa1-GFP5(S65T) | WT | Col-0 | N/A | HO | kan | (Dettmer et al.)<sup>45</sup> |
| SYP61-GFP | 35S::SYP61-GFP | WT | Col-0 | N/A | HO | N/A | (Drakakaki et al.)<sup>46</sup> |
| Rab A2a-YFP | Rab-A2a::YFP-Rab-A2a | WT | Col-0 | N/A | HO | N/A | (Chow et al.)<sup>47</sup> |
| Knolle-GFP | KNOLLE::GFP-KNOLLE | WT | Ler | N/A | HO | N/A | (Boutte et al.)<sup>48</sup> |
| VAMP711-Cherry | N/A | WT | N/A | N/A | HO | N/A | (Geldner et al.)<sup>49</sup> |
| SYP22::SYP22-YFP | SYP22::SYP22-YFP | WT | Col-0 | N/A | HO | N/A | (Nodzynsky et al.)<sup>50</sup> |
| gammaTIP-GFP | 35S::gammaTIP-GFP | WT | Col-0 | N/A | HO | kan | (Saito et al.)<sup>51</sup> |
| PEN1-GFP in pen1-1 KO background | 35S::GFP-PEN1 | pen1 KO | Col-0 | N/A | HO | N/A | (Nielssen et al.)<sup>52</sup> |
| DRP1a-GFP | 35S::DRP1a-GFP | WT | Col-0 | N/A | HO | kan | (Fujimoto et al.)<sup>53</sup> |
| CLC-GFP | CLC2::CLC-GFP | WT | N/A | N/A | HO | N/A | (Ito et al.)<sup>53</sup> |
| secGFP | 35S::Sec-GFP | WT | Col-0 | N/A | HO | hyg | (Zheng et al.)<sup>54</sup> |
| GFP-VSP35B in vps35b-2 KO | VPS35B::VPS35B-GFP | vps35b KO | Col-0 | N/A | HO | kan | (Munch et al.)<sup>55</sup> |

Abbreviations: BY-2 = bright yellow 2; TT = tandem tag; Fluc = firefly luciferase; Rluc = *Renilla* luciferase; dLuc = dual luciferase; BG = background; Acc = Accession; Gen = Generation; HO = Homozygous; PR = Plant resistance marker; WT = wildtype; KO = knockout; N/A = not available.

