## Supplementary Video 1 for "Chemical screening pipeline for identification of specific plant autophagy modulators"

### Slide 1
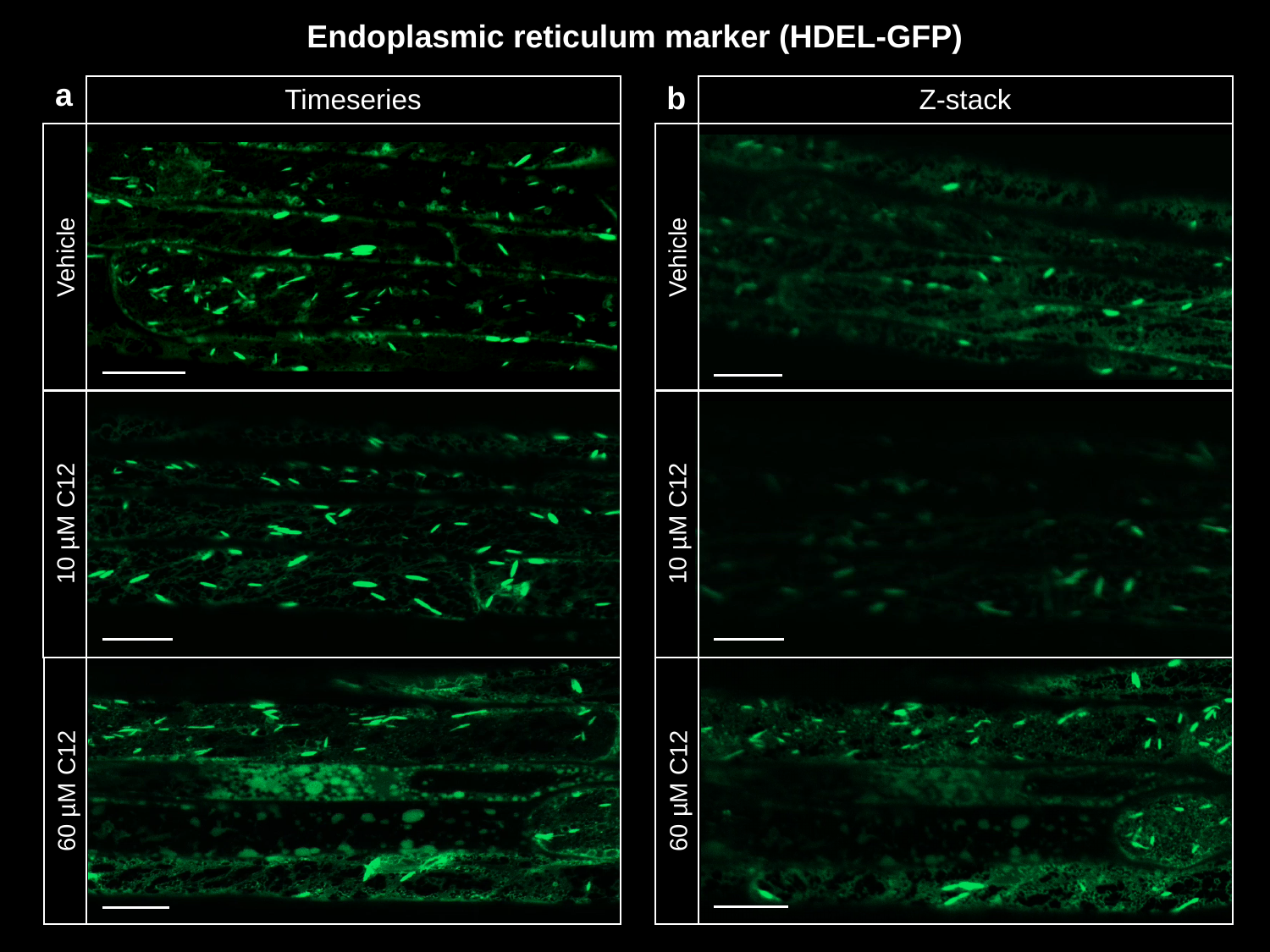

Endoplasmic reticulum marker (HDEL-GFP)
a
b
Timeseries
Vehicle
10 µM C12
60 µM C12
Z-stack
Vehicle
10 µM C12
60 µM C12
