## Supplementary File 1 for "Chemical screening pipeline for identification of specific plant autophagy modulators"

**Tandem tag assay optimized for semi-automated autophagy flux measurement in *Arabidopsis thaliana* roots**

**Abstract**

Autophagy is the major catabolic process in eukaryotes with a key role in cell homeostasis. *In vivo* measurement of autophagic activity is pivotal for investigating the role of the pathway in organism development and stress responses. Here we describe optimization of the tandem tag assay for detection of autophagic flux *in planta* in epidermal root cells of *Arabidopsis thaliana* seedlings. For this assay, ATG8 is fused to the tandem tag that consists of two fluorescent proteins, (TagRFP and mWasabi) and is expressed in wildtype or autophagy-deficient backgrounds to obtain reporter and control lines, respectively. Upon autophagy activation, the TagRFP-mWasabi-ATG8a fusion protein is incorporated into autophagosomes and delivered to the lytic vacuole. Ratiometric quantification of the lytic pH-tolerant TagRFP and pH-sensitive mWasabi fluorescence in the vacuoles of control and reporter lines allows for a reliable estimation autophagic activity. We describe a step by step protocol for plant growth, imaging and semi-automated data analysis. The protocol presents a rapid and robust method that can be applied for any studies requiring *in planta* quantification of autophagic flux.

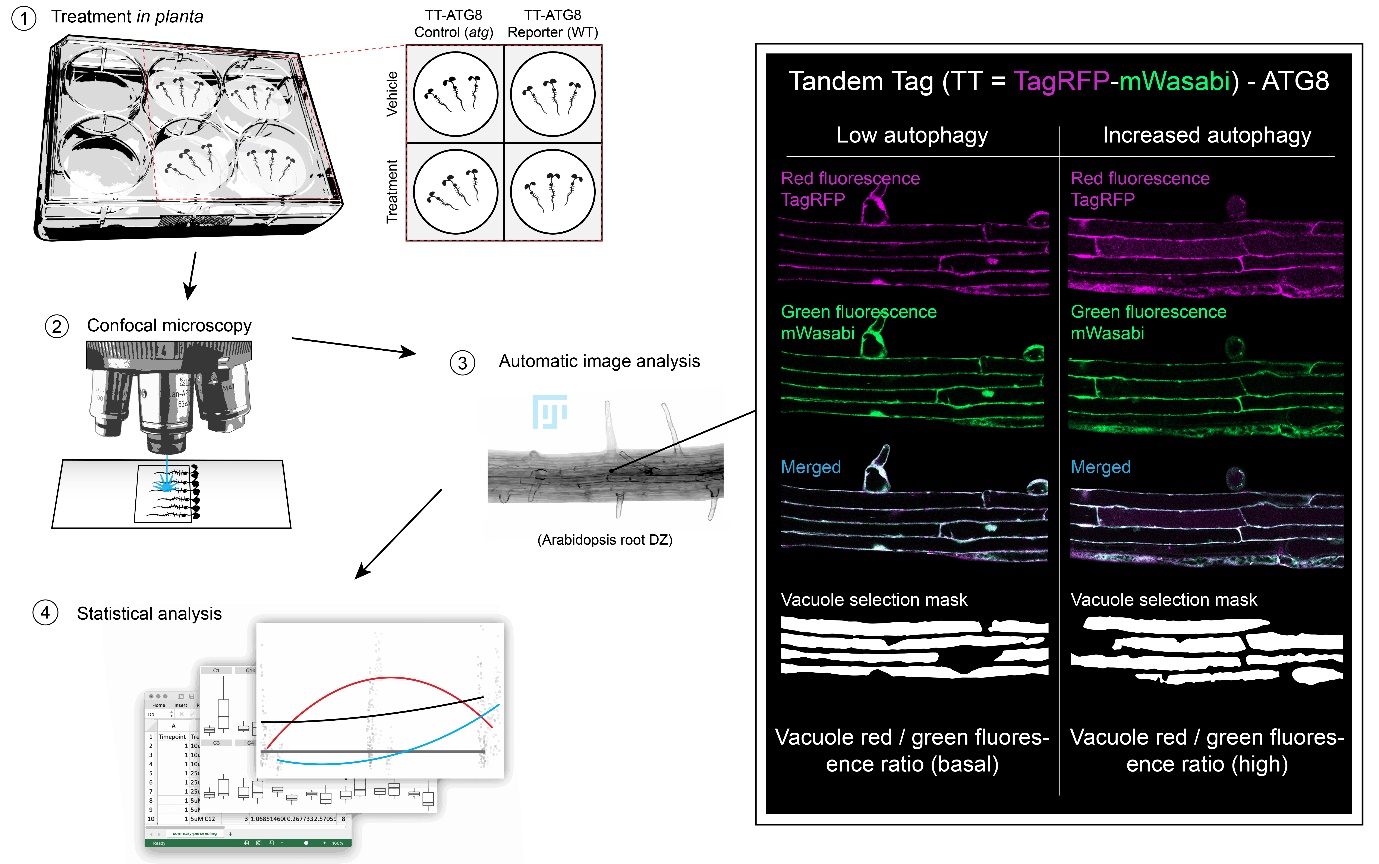

**Introduction**

The Tandem Tag (TT) assay employs ratiometric quantification of red and green fluorescence and is a widespread approach for quantifying autophagic flux in yeast and mammalian cells^1–3^. It has been previously described to be also applicable for plant cells in a study using tobacco BY-2 cell suspension cultures^4^. Another advantage of this assay is that it provides valuable morphological data including subcellular localization of ATG8 during puncta formation and translocation from the nuclei^5^. Here we provide a detailed protocol for *in planta* quantification of autophagic flux in epidermal root cells of *Arabidopsis thaliana* seedlings.

For this protocol we used stable transgenic Arabidopsis lines expressing the optimized Tandem Tag of TagRFP and mWasabi^3^ fused to AtATG8a and driven by a double 35S promoter. The TT-AtATG8a fusion was introduced into wild-type (WT) or autophagy-deficient (*atg5atg7*) backgrounds, to produce reporter and control lines, respectively. Under normal growth conditions TT-AtATG8a is localized in the cytoplasm and in the nuclei of the root epidermal cells, but upon induction of autophagy it is gradually translocated to the cytoplasm, where it is incorporated into autophagosome membranes and delivered together with the cargo to the lytic vacuole. While the red fluorescence of TagRFP shows relatively high tolerance to the low pH of the lytic compartments^3^, green fluorescence of mWasabi is significantly reduced under the same conditions. Thus, ratiometric measurement of the TagRFP and mWasabi fluorescence allows to estimate the delivery rate of the fusion protein to the lytic compartment and eliminates potential bias caused by the differences in fusion protein expression. Furthermore, since the assay relies on confocal microscopy image acquisition and analysis, it is possible to obtain time-resolved and dose-dependent data about the changes in autophagic activity. Although we describe only a protocol for detection of the TT-AtATG8a delivery to the vacuole, it is also possible to use the same imaging data to also visualize and quantify dynamics of autophagosome formation.

We observed that response to known modulators of autophagy activity, such as AZD8055 and Concanamycin A (ConA)^1^, in Arabidopsis roots significantly varied depending on the area of the root zone imaged. By comparing data for different root zones, we established that the most reproducible measurements could be obtained by analyzing images of root epidermal cells located in the beginning of the differentiation zone. Nevertheless, we still observed quite significant variation between responses in tricho- and atrichoblasts. Hence, a relatively large number of images was required for the data analysis to obtain a reliable mean value representative of average autophagy activity in the root zone.

To facilitate the measurements, we developed a semi-automated high-throughput image analysis protocol described below. It relies on the use of macros written in ImageJ Macro Language (IJL) and R scripts. The protocol was tested using images obtained with Zeiss and Leica confocal laser scanning microscopes (CLSMs) and should be applicable for data from other manufacturers of CLSMs. Furthermore, to enable applicability of the protocol for images obtained on different CLSM systems that will naturally vary in efficacy of detection, we added an extra step of threshold values adjustments that will maximize the capacity of the vacuole area selection tool.

Importantly, the protocol described here allows quantification of an autophagy-dependent delivery rate of ATG8 to the lytic vacuole, which is a good proxy for the level of autophagic activity. However, in order to make sure that autophagic flux is completed, it is recommended to combine this protocol with other assays verifying degradation of autophagic cargo e.g. the GFP-ATG8 cleavage assay or long-lived proteins assay^6^.

**Reagents and biological material used**

1. Bleach solution: 10% Klorin (12.5% sodium hypochlorite, Sweden), 0.05% Tween-20.
2. 0.5x MS medium liquid medium: 0.5x Murashige and Skoog medium with vitamins (M0222, Duchefa), 10 mM MES (M8250, Sigma), 1% sucrose (S0809, Duchefa), pH 5.8. Sterilized by autoclaving.
3. 0.5x MS solidified medium: 0.5x MS medium as described above solidified with 0.8% Plant agar (P1001, Duchefa)
4. DMSO (or other vehicle)
5. AZD8055 (S1555, Selleckchem)
6. Concanamycin A (ConA; C9705, Sigma)
7. Immersion suitable for the CLSM used for imaging
8. Reporter line AD51: *Arabidopsis thaliana*, accession Col-0, homozygous line expressing TagRFP-mWasabi-AtATG8a under control of 2x35S promoter in the wild-type background (reference to the methods paper Dauphinee et al., 2019)
9. Control line AD4: *Arabidopsis thaliana*, accession Col-0, homozygous line expressing TagRFP-mWasabi-AtATG8a under control of 2x35S promoter in the *atg5-1/atg7-2* double knockout background (reference to the methods paper Dauphinee et al., 2019)

**Equipment**

1. 92 x 16 mm, PS, with ventilation cams, Petri dishes (82.1473.001, Sarstedt) or 120x120x17 mm square PetridDishes with vents (688161, Greiner Bio-One)
2. Soft tweezers (FST, Dumont Biology)
3. Pipettes for 100-1000 µl and 1-10 μl
4. 6-well tissue culture plates (83.3920, Sarstedt)
5. 360° Vertical Mini Rotator (PTR-25, Grant-bio)
6. 4 °C fridge
7. A growth cabinet/growth room suitable for *Arabidopsis thaliana*: 20-22°C, 50-70 % humidity, 150 µM light (full light spectrum)
8. Microscope slides (631-1550, VWR)
9. Cover slips 25 x 75 mm (11911998, Fisher Scientific) and 25 x 25 mm (15707593, Fisher Scientific)
10. Confocal Laser Scanning Microscope (CLSM; LSM 800, Zeiss). The protocol was verified on the data obtained using Leica SP5 and Leica SP8 CLSM.

**Procedure**

1. Seed sterilization (40 min)
   - Place ca 15 μl of seeds of reporter and control lines into 1.5 ml Eppendorf tubes.
   - Add 1.5 ml of the bleach solution.
   - Incubate the seeds for 30 min, agitating with a 360° vertical rotator.
   - Under sterile conditions pipette out the bleach solution.
   - Add sterile water to the seeds. Mix and pipette it out.
   - Perform the wash with sterile water at least three times to wash out the leftovers of the bleach solution.
   - Seeds can be vernalized either in the sterile water or as described in the step three.

2. Seed plating (40 min)
   - Use 1 ml pipette to transfer individual seeds in rows onto a Petri dish with solid 0.5x MS medium. Keep ca 2–5 mm distance between the seeds in the same row to avoid entanglement of roots. Keep ca 5 cm vertical distance between the rows (Fig. 1a).
   - Seal the plates with a strip of Parafilm or a strip of Saran wrap.

3. Vernalization (24–48 h)

- Incubate the plates in the dark at 4°C for 24–48 h

4. Growth on vertical plates (5d)
   - Transfer the plates to the growth cabinet (150 µM light, 22°C for 16 h, darkness, 20°C for 8 h). Make sure the plates are perfectly vertical to enable root growth on the top of the medium (Fig. 1b).
   - Grow the seedlings on the plates for approximately 5 d, the root length should reach 2 cm.

5. Drug treatment (2-24h)
   - Pipette 3 ml of liquid 0.5x MS medium into wells of a 6-well tissue culture plate. Add the chemical compounds of the required concentration.
   - Using soft tweezers gently pick seedlings from the plates and transfer into liquid 0.5x MS medium (maximum 20 seedlings/well)
   - Gently pipette the medium from the well onto the seedlings in the well to submerge the roots (white arrow, Fig. 1c).
   - Seal the plate with a strip of Parafilm or Saran wrap.
   - Incubate the plate under the same growth conditions for the required amount of time.
6. Mounting the samples (1 min)
   - Pipette ca 50 μl of medium from a well on the 25 x 75mm cover slip.
   - Using soft tweezers, gently pick seedlings from the well and place them onto a drop making sure that the root is straight.
   - Place up to six seedlings on the cover slip.
   - Cover the roots with 25 x 25 mm cover slip (Fig. 1d)
   - Apply immersion medium if needed to the objective lens and place the sample on the microscope stage.
7. Setting up scanning parameters for confocal laser microscopy (30 min, required only once)
   - To estimate optimal range of settings for scanning, it is advisable to perform a pilot experiment using three control treatments with 500 nM AZD 8055 for 4 h, 500 nM ConA for 6h and a corresponding treatment with vehicle (usually 0.05% DMSO).
   - Configure settings for sequential scanning in two channels, please note that the channel order is important for further analysis:
     - Channel 1 for detection of mWasabi: excitation at 488 nm, emission detection range 490–564 nm.
     - Channel 2 for detection of TagRFP: excitation at 561 nm, emission detection range 564–700 nm.
     - It is advisable to use the most sensitive detectors available in the system (i.e. GaAsp or HyD detectors for Zeiss or Leica CLSM, respectively).
     - Using 40x objective is advisable to obtain images most applicable for automated analysis, as this magnification provides high quality of imaging even for low signal and still allows a relatively large field of view.
     - Set switching between channels for each frame to minimize the crosstalk between channels and optimize the pinhole size. If possible, use the pinhole of 1 AU for each of the channels.
     - If possible, use at least 16-bit resolution to increase the resolution of intensities.
     - The sample treated with AZD 8055 will have the weakest fluorescence and should be used to adjust laser intensity and the Detector Gain (Master Gain) to the lowest possible values that still provide detectable signal.
     - The sample treated with ConA will have the strongest fluorescence at least in the green channel and should be used to re-adjust laser intensity and the Detector Gain (Master Gain) to the lowest possible values that do not result in oversaturated pixels.
     - The sample treated with the DMSO can be used to verify the applicability of the adjusted settings.
     - Additionally, noise can be decreased by ramping down the scanning speed or increasing the averaging number. To our experience, scanning at the speed of ca 10 sec per frame produced images of the quality appropriate for further analysis and also resulted in acceptable time for experiment.
8. Scanning (ca 30 min per treatment)
   - Most of CLSM software will have an option of scanning at selected positions that will significantly decrease the time required for the experiment.
   - Mark the positions at the beginning of the differentiation zone of the root (Fig. 1e).
   - Start fast scanning mode (live scan) and readjust the focal plane for each position to the middle section through the vacuole of the epidermal cells. Good, bad and ugly quality scans will greatly affect the vacuole selection (Figs. 1f, g).
   - When all positions are readjusted, acquire and save the images. Please note, that automated statistical analysis will use information provided in the names of files. Please use the following rules to introduce the required parameters into the name of your images:
     - Separate parameters by underscores (_)
     - Name the files as following: ***line***_***treatment***_seedling***X***_image***Y*** (e.g. *Reporter_50uM.C12_seedling1_image2.czi*)

Where *line* and *treatment* are text variables or strings identifying the lines (either “Reporter” or “Control”) and treatments (i.e. “50uM.C12”) used. One treatment must be named “Vehicle”. *X* and *Y* are numbers indicating seedling and image replicates thus representing biological and technical replicates, respectively. Note that “seedling” and “image” are fixed strings and only the numbers *X* and *Y* are changed from image to image.

9. Data processing (ca 10 minutes or ca 40 minutes including installation of the required software)

Data processing is done in four steps: (i) confocal images are converted into .TIF files; (ii) adjustment of the threshold values to optimize recognition of the vacuoles must be performed while using the protocol for the first time and, if needed, can be redone for individual experiments; (iii) fluorescence intensities of the vacuoles are quantified and saved; (iv) quantification of the ratios and plotting of the data as ratios in control *vs* reporter lines, or ratios *vs* time and concentration.

- Place all images for processing into a single folder. If several experiments should be processed, place them as subfolders into a single folder.
- Download and install Fiji, the version of ImageJ with included set of plugins (<https://fiji.sc/>; for this study, we used versions 1.51s and 2.0.0-rc-69/1.52i).
- Download and install R^7^ (https://www.r-project.org, we used 3.5.2 and 3.5.1) and RStudio (<https://www.rstudio.com/>; we used versions 1.1.453 and 1.2.1186). The first time you use R you need to make sure its dependencies are installed. Either use RStudio’s interface for installing packages, and install the packages ggplot2^8^, dplyr^9^, and readr^10^, or install them manually by typing the following into the console:

install.packages(c("ggplot2", "dplyr", "readr"))

- Download the GitHub repository as a zip file (<https://github.com/jonasoh/AuTToFlux>). Alternatively, RStudio can be used to automate repository cloning: firstly, make sure that GIT is installed (<https://git-scm.com/downloads>); then proceed in RStudio with starting a new project (File -> New Project -> Version Control -> GIT and input the following URL: <https://github.com/jonasoh/AuTToFlux>. The benefit of cloning the repository in RStudio is the possibility of obtaining updates *via* the "pull branches” option (Tools -> Version Control -> Pull Branches.
- Run the Image Processor: launch Fiji/ImageJ and go to Plugins -> Macros -> Run -> locate ImageProcessor.ijm.
- The image processor macro opens all compatible images in the target folder and saves them as .TIF files; original acquisition dates of the images are saved in separate files (*.time). Multi-image files from experiments acquired using the “Positions“ function are split so that each image is saved as a separate .TIF file.
- Optimize the threshold parameters for selecting the vacuoles:
  - Copy at least three .TIF images generated by the ImageProcessor into a separate folder. Aim to select images representing the best, the worst, and average quality.
  - Launch Fiji/ImageJ and go to Plugins -> Macros -> Run -> locate CalibrateThreshold.ijm. This runs the macro. In the file picker immediately presented, locate the folder with representative images. Macro will record the area sizes corresponding to the vacuoles and also save masks as tif files containing threshold values in their names, e.g. a file named *.thr0-3.TIF would correspond to the mask created with the threshold values (0;3). The results overview will be generated as threshold-overview.TIF (Fig. 1g).
  - Launch RStudio.
  - Open the RStudio project file (AuTToFlux.Rproj).
  - To estimate what threshold value provides the largest selected vacuole area for all analyzed images, run the EvaluateCalibration.R script (select the R script, click Code -> Source and then choose the folder containing the .CSV files generated in the previous step – Note: on non-Windows systems you need to type in the pathname of the folder).
  - The script will plot average sum vacuole areas selected on all analyzed images *vs* threshold values. Select the threshold value that corresponds to the largest area (Fig. 1h).
  - Using the information gathered from the threshold-overview and the threshold graph, select an appropriate upper threshold value for masking all images using the FluorescenceIntensity macro as described below. The masks should not contain any background areas.
  - Before proceeding with image thresholding and data analysis, ensure that the calibration folder is removed from the directory containing the processed images.

- Run the threshold macro: launch ImageJ -> Plugins -> Macros -> Run -> locate and open the macro file FluorescenceIntensity.ijm. The macro will prompt you to select the threshold value determined earlier in the calibration process. For each image the threshold macro automatically selects areas corresponding to the vacuoles using the green (mWasabi) channel and saves the masks for the selected areas as .TIF files. The mask is then used to quantify intensities of red and green fluorescence in the corresponding channels of the image. Ratios of vacuolar red/green fluorescence intensities are saved to .CSV files with matching filenames.

10. Data analysis

- Launch RStudio.
- Locate the R project file in the directory created from the GitHub repository and open it.
- To estimate dose and time-dependent responses the red/green (TagRFP/mWasabi) ratios are plotted *vs* time using Flux-vs-Time.R (select the R script, click Code -> Source and then choose the folder containing the .CSV files generated in step 9).
  - The initial time of treatment (format YYYY-MM-DD HH:MM, 24-h time) is user-specified by creating a tab-delimited text file (using any spreadsheet software, e.g. Microsoft Excel) with two columns (named "Treatment" and StartTime"), which should be saved as info.txt in the folder containing the .CSV files for analysis.

e.g.

| Treatment | StartTime |
| --- | --- |
| Vehicle | 2018-07-06 09:50 |
| Treatment | 2018-07-06 10:00 |

- A summary of the experimental data is generated with ratios normalized to the vehicle and grouped by treatment and seedling number (Fig. 1i). To plot the data, we recommend ggplot2 with settings: axes x=Elapsed time, y=Normalized ratios, and color=Treatment.
- Further statistical analysis will depend on the treatments present in the experiment and can be performed using R or other software, e.g. Origin (<https://www.originlab.com/>) or JMP (<https://www.jmp.com/>).
- The data can be also used to estimate IC50 using R or other software, e.g. Prism (<https://www.graphpad.com/>).
- To demonstrate autophagy-dependent response, red/green ratios of control lines are plotted *vs* ratios of reporter lines. For this, run Control-vs-Reporter.R (select the R script and then click RStudio -> Code -> Source) and then choose the folder containing the .CSV files generated in step 9).
  - This script may take as its argument either a folder containing several treatments as subfolders, or a folder without subfolders that contains a single experiment.
  - The script summarizes red/green ratios (normalized to vehicle) and groups by line (control or reporter), treatment and seedling number. An unpaired, two-tailed Student’s t-test is used to compare log-transformed normalized means of the control and reporter groups. A summary table named pvals.txt is generated. If p-values derived from permutation (i.e. exact p-values) are desired, uncomment the appropriate sections marked in the R script.

Note: for both Control-vs-Reporter.R and Flux-vs-Time.R the data is saved both as raw data (as summary-full.txt in the experiment directory) and as per-seedling summary statistics (summary-perseedling.txt), to facilitate analysis using other statistical software.

**Troubleshooting**

For additional troubleshooting information please refer to Table 1.

**Conflicting financial interests**

The authors declare no competing financial interests.

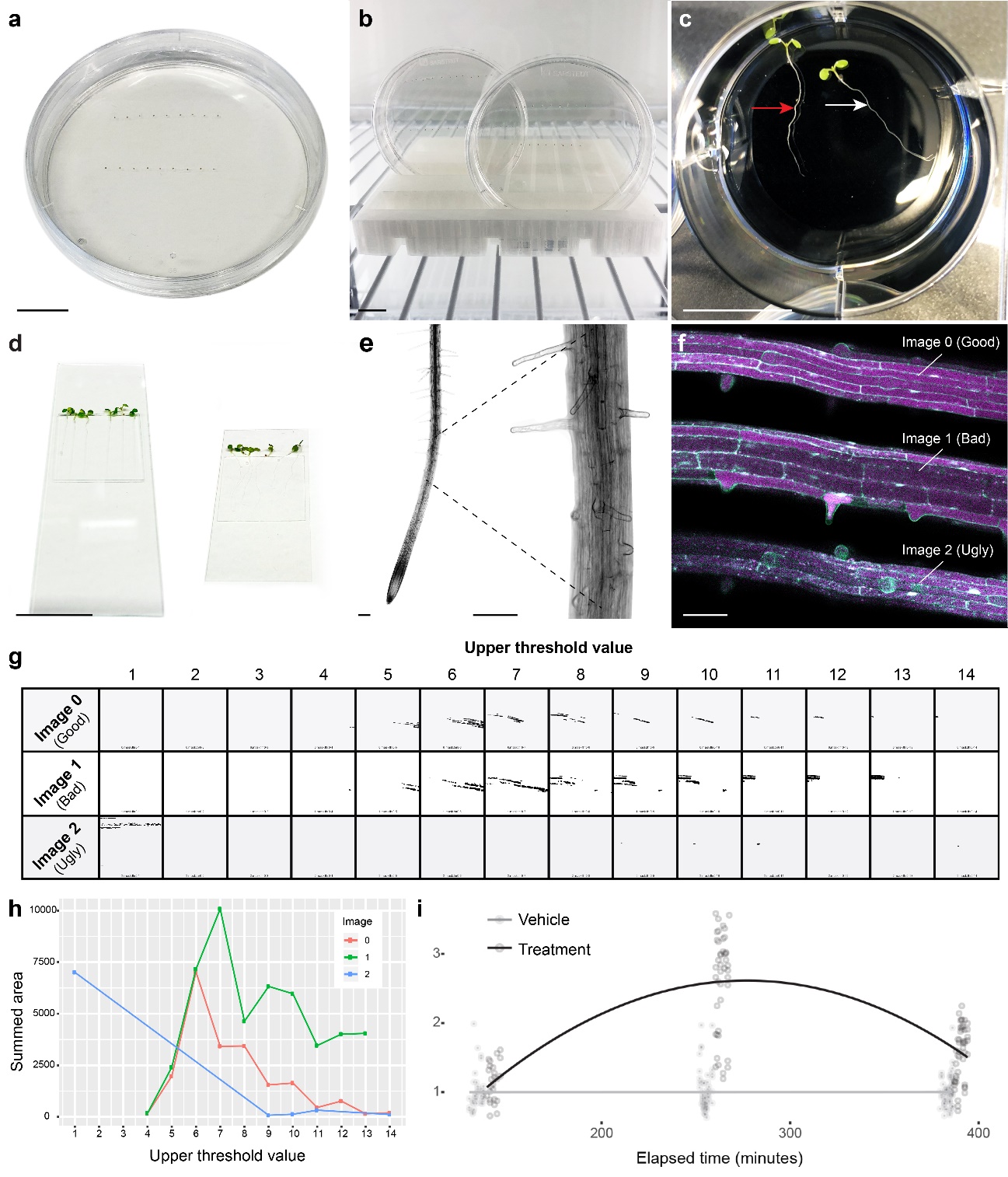

**Fig. 1: Arabidopsis tandem tag autophagic flux assay.**

**a,** Tandem tag (TT; TagRFP-mWasabi-ATG8a) reporter (TT in WT background) and control (TT in autophagy-deficient background) *Arabidopsis thaliana* seeds plated on 0.5x MS medium and grown vertically (**b**)**. c,** *Arabidopsis* seedlings placed in a 6-well plate for treatment. Note that the roots should be submerged gently (white arrow) and should not be floating on the surface (red arrow). **d,** Seedlings mounted on a glass slide (left) or coverslip (right) for CLSM. **e,** Epidermal layer of the root differentiation zone **f,** Images 0, 1 and 2 represent good, bad and ugly quality scans of differentiation zone vacuoles in the reporter line. **g,** Vacuole layer masks generated during calibration using the CalibrationThreshold.jim macro. Note that ugly scans (**f**) may lead to complete or partial failure of vacuole selection. **h,** Sum vacuole areas selected on all analyzed images (3 recommended) *vs* threshold values generated by the EvaluateCalibration.R script. **i,** Flux-vs-Time.R script output for vehicle and a single concentration for one treatment generated in R using ggplot2. Scale bars, 1.5 cm (**a-d**); 50 µm (**e, f**).

**Table 1. Troubleshooting information for the tandem tag assay**

| **Problem** | **Possible cause** | **Solution** |
| --- | --- | --- |
| Low number of data points | High noise on the images | 1. Adjust scanning settings to decrease the noise 2. Adjust threshold values during data analysis |
|  | Focusing not on the middle section of the vacuoles. | Make sure that the roots on the samples are not drifting during imaging and are placed in the thinnest possible layer of solution. |
|  | Wrong threshold values in the FluorescenceIntensity macro | Follow the instructions in the procedure for estimating optimal values and adjusting the macro. |
| Incorrectly assigned groups in Flux-v-Time.R | Different number of data points for each timepoint | The script requires the number of data points for each group to be roughly equal. Remove or add data points to balance the experiment. |
| Time-v-Flux.R exits with error "Error in eval(ei, envir) : file.exists(infofile) is not TRUE" | There is no info.txt in the directory. | Create the info.txt file according to the instructions. |
| Time-v-Flux.R gives unexpected results | Times in info.txt do not match the acquisition times of the images. | Make sure that times in info.txt correspond to times as recorded by the microscope. |
|  | Treatments in info.txt do not match filenames. | Make sure that treatment names are given exactly the same in filenames and info.txt. |
| Data points are duplicated | ImageProcessor has been run twice on the same set of images. | Delete all .TIF files in the directory and rerun ImageProcessor on the files directly from the microscope. |
| Script exists with error "*file* does not match naming scheme". | File is incorrectly named. | Double check that the filename matches the naming scheme described in the protocol. |
